# Complex interaction of resource availability, life-history and demography determines the dynamics and stability of stage-structured populations

**DOI:** 10.1101/138446

**Authors:** Sudipta Tung, M. Rajamani, Amitabh Joshi, Sutirth Dey

**Affiliations:** Population Biology Laboratory, Biology Division, Indian Institute of Science Education and Research-Pune, Dr. Homi Bhabha Road, Pune, India, 411 008; Evolutionary Biology Laboratory, Evolutionary and Organismal Biology Unit, Jawaharlal Nehru Centre for Advanced Scientific Research, Jakkur P.O., Bengaluru, India, 560 064; Clintus Network Ltd., No. 205/1 & 2, PCR Warehouse Complex, Soukya Road, Koralur Village, Hoskote, Bengaluru 562 114

**Keywords:** fluctuation index, stability, constancy, persistence, minimum critical size, time-series, stage-structured model, Sterile Insect Technique

## Abstract

The dynamics of stage-structured populations facing variability in resource availability and/or demographic factors like unequal sex-ratios, remains poorly understood. We addressed these issues using a stage-structured individual-based model that incorporates life-history parameters common to many holometabolous insects. The model was calibrated using time series data from a 49-generation experiment on laboratory populations of *Drosophila melanogaster*, subjected to four different combinations of larval and adult nutritional levels. The model was able to capture multiple qualitative and quantitative aspects of the empirical time series across three independent studies. We then simulated the model to explore the interaction of various life-history parameters and nutritional levels in determining population stability. In all nutritional regimes, stability of the populations was reduced upon increasing egg-hatchability, critical mass and proportion of body resource allocated to female fecundity. However, the stability-effects of increasing sensitivity of female-fecundity to adult density varied across nutrition regimes. The effects of unequal sex-ratio and sex-specific culling were greatly influenced by fecundity but not by levels of juvenile nutrition. Finally, we investigated the implications of some of these insights on the efficiency of the widely-used pest control method, Sterile Insect Technique (SIT). We show that increasing the amount of juvenile food had no effects on SIT efficiency when the density-independent fecundity is low, but reduces SIT efficiency when the density-independent fecundity is high.

## 1. INTRODUCTION

The quality and quantity of available nutritional resources affect the growth, development, and overall survival of organisms. This, in turn, can have a major impact on population stability and dynamics (Cole and Batzli 1979; Turchin and Batzli 2001). Rapid global climate change and habitat degradation due to human activity, has increased the spatial and temporal variability of food available in many parts of the world (Walther et al. 2002). That is why, understanding how the interaction of nutritional availability with different life-history traits and demographic factors shapes the dynamics of populations, has become an important problem in modern population ecology (Oro et al. 2004; Turchin and Batzli 2001).

Over the last two decades, a number of studies on vertebrate species, have yielded valuable insights on this topic (reviewed in Boutin 1990). For example, human-induced factors, like industrial fisheries and refuse dumps, and the resulting increase in food supply, has led to a substantial increase in the population size of several opportunistic seabird species (Croxall and Rothery 1991; Garthe et al. 1996). On the other hand, food-related stress at the time of reproduction can have complex effects on population processes in some seabird species (Kitaysky et al. 2010), which can subsequently affect their abundance (Byrd et al. 2008; Oro et al. 2004). Similarly, a theoretical study on the population dynamics of arvicoline rodents (Turchin and Batzli 2001) have shown that food limitation plays an essential role in maintaining the periodic fluctuations observed in this system.

In terms of species diversity, invertebrates in general and insects in particular, far outnumber the vertebrates (Stork 1997). Insects also play important roles in maintaining the health of ecosystems (Yang and Gratton 2014) and have considerable economic importance (Losey and Vaughan 2006). Consequently, the effects of nutritional availability on dynamics of insect species has also been an area of interest for ecologists (Abbott and Dwyer 2007). Unfortunately, it is not clear how much of the insights gathered from the vertebrate systems mentioned in the last paragraph, are also applicable to invertebrates. One of the reasons for this is the fact that most insect life-cycles are stage-structured, which means different developmental stages can potentially have different kinds of nutritional requirements (Boggs 1981). Not surprisingly, the dynamics of an insect population can be a fine interplay between the quality and quantity of resources available to the various developmental stages (Mueller 1988). Unfortunately, in spite of its considerable scientific and practical implication, this aspect has remained relatively less explored among invertebrate populations (although see Costantino and Desharnais 2012).

The situation becomes even more complex for sexually reproducing species, when nutritional availability affects and/or interacts with demographic factors, such as sex ratio, which can have strong impact on the overall dynamics (Johnson 1994). Simple models of population dynamics often assume that the entire dynamics can be understood in terms of the number of females in the population (May 1976). Over the last few decades, the consequences of relaxing this assumption has been extensively investigated both theoretically and experimentally (reviewed in Rankin and Kokko 2007). This line of study helps us understand the dynamics of those populations in which the two sexes face very different rates of mortality due to natural or anthropogenic reasons (Berec et al. 2016; Halvorsen et al. 2017; Perlman et al. 2015). Moreover, one of the highly successful pest eradication strategies, the Sterile Insect Technique (SIT), explicitly depends on altering the sex ratio of the population (Dyck et al. 2005). In nature, the amount of food available would obviously vary across space and time and this, in principle, can modulate the effects of sex ratio on the dynamics. Yet, the interaction of food and sex ratio in shaping the dynamics in general and stability in particular, has remained relatively poorly understood.

Investigating these issues is no easy task. This is because, to begin with, one needs an extremely thorough knowledge of how the various determinants of the dynamics (like fecundity, survivorship etc.) are affected by the nutrition available to each developmental stage. Then, these insights have to be combined to predict how the dynamics of the population are affected by the interactions. Then one needs to factor in demographic changes, like altered sex-ratio, to understand how they interact with the various nutrition levels to shape the dynamics. The complexity of the problem makes it difficult to conduct a completely empirical study even on laboratory populations, to say nothing about natural ones. On the other hand, since a large number of interactions are involved, a completely theoretical study, based on modelling from first principles, can be difficult to interpret. This is because, any modelling study, by its very nature, forces to the modeler to make assumptions about the nature of the real system at every step. Thus, approximation errors and interactions can compound and lead to results that might be hard to interpret.

To address this issue, we used a multi-stage strategy. We chose the fruitfly, *Drosophila melanogaster* as our model system, primarily because the laboratory ecology of this species, particularly in the context of nutrition, has been well worked out (reviewed in Mueller 1985; Mueller and Joshi 2000). Using these empirical insights, we first developed a stochastic, stage-structured model of *Drosophila* population dynamics. The model incorporated life-history parameters like hatchability, critical mass, adult body size, and fecundity, which are common to many holometabolous insect populations. Then, in order to calibrate the model parameters and validate the model outcomes, we conducted a 49-generation long experiment using laboratory populations of *D. melanogaster* subjected to four different nutritional regimes. We then compared the experimental data with our simulation results to show that our model is able to capture various qualitative and quantitative aspects of *Drosophila* population dynamics in the laboratory. We further demonstrated the performance of our model for understanding the dynamics of *D. melanogaster* populations in three ways. First, we used it to resolve a discrepancy between observations from an earlier study and our results. Second, we used it to generate clear predictions about how various life-history-related parameters affect the dynamics of the populations under the various nutritional regimes. Third, we used data from a previous experimental study to validate some of these predictions.

After thoroughly establishing the bona-fides of our model as a descriptor of *Drosophila* dynamics, we briefly compared its various features and predictions with those arising from models and experiments on other species. We then used our model to investigate the various facets of the effects of unequal numbers of males and females on population dynamics. Our simulations suggested that this interaction is best understood in conjunction with the density-independent fecundity of the organisms. In a nutshell, the effects of changing the sex ratio on population stability and average size are dependent on certain aspects of the provided nutrition and not others. However, when it comes to SIT, the efficacy of the program depends more critically on the food provided over a large and biologically relevant parameter range.

## 2. MATERIALS AND METHODS

### 2.1 Experiment

The complete details of this experiment have been reported in the PhD thesis of one of the authors (Dey 2007). Here we present a brief account of the salient points and refer the readers to the SOM (Text S2) for more details.

The experiment consisted of thirty-two populations of *D. melanogaster*, each represented by a single vial culture. These 32 populations were randomly allotted to one of four nutritional regimes, such that there were eight populations per regime. Following established norms (Mueller and Huynh 1994; Mueller et al. 2000) these regimes were called HH, HL, LH, LL — where the first letter indicates the quantity of larval food and the second letter represents the status of adult nutrition. In case of larval food, H and L denoted ∼6 mL and ∼2 mL of banana-jaggery medium per vial, respectively, whereas in the case of adult nutrition, H and L referred, respectively, to the presence and absence of live yeast paste supplement to banana-jaggery medium. Thus, for example, HL denotes a nutritional regime comprising of ∼6 mL medium per vial for the larvae, but no live yeast paste supplement for the adults, and so on. The experiment was started with 8 adults (4 males and 4 females) in each population in the first generation, and the number of adults in each successive generation were noted. The experiment was continued for 49 discrete generations and yielded 32 time-series (see Figure S3 for representative time series).

### 2.2 Statistical analyses

Distributional properties of the experimental time series of adult populations were assessed using the mean, median, 5^th^, 10^th^, 25^th^, 75^th^, 90^th^ and 95^th^ percentiles in box plots (Zar 1999). The constancy stability (Grimm and Wissel 1997) of the populations was measured as the fluctuation index (*FI*, Dey and Joshi 2006) which is given as:

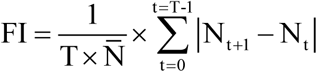

where 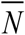 is the mean population size over *T* generations and *N_t+1_* and *N_t_* are the population sizes at the (*t+1*)^th^ and *t*^th^ generations, respectively. This formulation suggests that *FI* increases when the variance of the time series is increased, or the mean or the strength of autocorrelation of the time series is decreased.

In order to investigate the effects and interaction of larval and adult nutritional levels on constancy stability, the *FI* data were subjected to a two-way analysis of variance (ANOVA) with larval nutrition (fixed factor, two levels: Low and High) crossed with adult nutrition (fixed factor, two levels: Low and High). All statistical analyses were performed using STATISTICA^®^ v5 (StatSoft. Inc., Tulsa, Oklahoma).

### 2.3 The model and simulations

#### 2.3.1 Model formulation

Our individual-based, stochastic model (see S3 for the details of model formulation) is based on a previous deterministic model (Mueller 1988) with a few modifications (see section S5). The biological justifications for the various functional forms used and the model-calibration process has been described in detail in the SOM (see Text S4). Here we present a brief account to highlight the salient features of the model. A schematic diagram of the model processes has been included as Figure S2 for visualizing the sequence in which model processes were executed. Terms in italics denote parameters in the model, whereas state variables are in the `Courier New` font. For a given generation, the model starts with the number of eggs (`numegg`), a fixed fraction (*hatchability*) of which hatch to form `numlarva`, i.e. number of larvae:

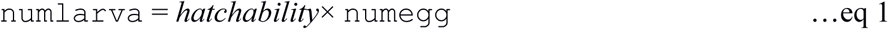

Each larva is then assigned a body-size (in arbitrary units of length) through a random draw from a normal distribution whose mean (mean_size) is positively related to the amount of food (*food*) in the culture but negatively related to `numlarva`:

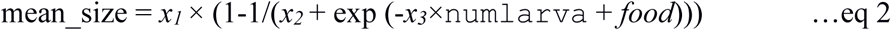

where *x*_1_, *x*_2_ and *x*_3_ are parameters (see Table 1 for descriptions) and exp denotes the exponential function. The standard deviation of the normal distribution is assumed to be a constant (=0.45).

**Table 1:** List of parameters used in the model.

| Parameter | Description | Value |
| --- | --- | --- |
| *food* | Amount of larval food present | 1.76 (LL and LH) and 2.56 (HL and HH) |
| *adnut* | Quality of adult nutrition/fecundity booster | 1 (LL and HL), 1.29 (HH) and 1.49 (LH) |
| *hatchability* | Egg-to-larval viability | 0.98 |
| $m_c$ | Critical mass *i.e.* the minimum mass/size required to become a viable adult | 1.1 (JB) and 1 (FEJ) |
| *sen_adden* | The coefficient of sensitivity of female-fecundity to adult density | 0.17 |
| *sen_adsize* | The coefficient of sensitivity of female-fecundity to adult size | 1.7 |
| *sigma_size* | Standard deviation of larval body size distribution | 0.45 |
| $x_1$ | Parameter | 2.5 |
| $x_2$ | Parameter | 1 |
| $x_3$ | Parameter | 0.009 |
| $x_4$ | Parameter | 1 |
| $x_5$ | Parameter | 85 |
| $x_6$ | Parameter | 2 |

Larvae which are larger than a critical minimum size (*m_c_*) form the adults of the generation, while the remaining larvae are assumed dead. The adults are randomly assigned to be males or females and it is assumed that all females are fertilized irrespective of the number of males in the population (this assumption is relaxed in the simulations discussed in section 3.9). In *Drosophila*, as in many insects, fecundity of females is known to vary positively with body size (Honěk 1993). We modeled this positive relationship using a parameter for sensitivity of fecundity to adult body size (*sen_adsize*). Another factor that affects the density-independent fecundity of adult flies is the presence of protein supplement in the form of live yeast paste which boosts the fecundity of the female flies. This effect was modeled through an adult nutrition parameter called *adnut* such that a value of *adnut* =1 would represent the fecundity of the flies in the absence of yeast paste, while *adnut* >1 represented the fecundity-boosting effects of the yeast paste (see Text S4 for model calibration). Taken together, the adult density-independent fecundity of the i^th^ female was given as:

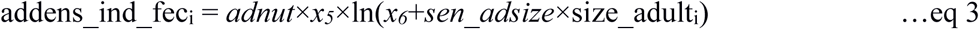

where size_adult_i_ is the size of the i^th^ adult and *x*_5_ and *x*_6_ are parameters.

In *Drosophila*, adult fecundity is negatively affected by adult density (Mueller and Huynh 1994; Rodriguez 1989). We modeled this effect using a hyperbolic function

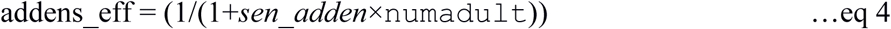

where the parameter *sen_adden* determines the sensitivity of female-fecundity to adult density (`numadult`). The number of eggs laid by each female is computed as the product of eq 3 and eq 4, and summed up over all females to form the `numegg` for the next generation.

#### 2.3.2 Simulations

To investigate the population size distributions for each nutritional regime (LH, HH, HH or HL), we simulated eight replicate runs of the model with 49 generations in each replicate, to keep parity with the experimental data. However, none of our conclusions changed when the length of the time series was increased (see Text S9 for discussion). Every simulation run started with 18 eggs. When there was extinction in any generation (i.e. `numadult` = 0), the time series was reset with four females with body size =2×*m_c_*. Following previous studies (Sah et al. 2013; Tung et al. 2014), we also incorporated additional demographic stochasticity in the model by considering a 50% chance of extinction whenever population size went below eight. If extinction occurred due to demographic stochasticity, the population was reset in the same way as mentioned above.

We also explored the effects of wide ranges of life-history related parameters (*hatchability, m_c_, sen_adden, sen_adsize*) on population stability. For each value of a given parameter, we took an arithmetic mean of *FI* measured over 100 replicate time-series, each of which was 100-generations long. All other conditions of the simulations were identical to those in the previous paragraph. Note that the larger number of longer runs in these simulations are appropriate as the goal of this part is to characterize the ‘theoretical’ effects of changes in those parameters, rather than comparing results to experimental data.

Our empirical data revealed that the HL regime had a greater average population size and lower *FI* compared to the HH populations (see section 3.5 for details) whereas an earlier study (Mueller and Huynh 1994) had shown that the population size of HH was greater than that of HL and the two regimes had similar constancy stability. In order to investigate this discrepancy between the two results, we simulated our model with five different values of larval food ranging from 3.0 to 7.0 in step size of 1.0. Each value of larval food level (food) was crossed with two values of adnut – 1.0 and 1.29-which represented the presence and absence respectively of yeast for the adults. For each *food × adnut* combination, we simulated eight 49-generation long time-series, and obtained the corresponding population size distributions, and *FI* and egg-to-adult viability values. All other parameter values were identical to the earlier simulations (Table 1).

### 2.4 36-generation simulation and experiment

To validate one of the predictions arising from our model, we compared our model output with the dynamics of four *Drosophila* populations selected for faster development and early reproduction (henceforth called FEJ_1-4_) for ∼125 generations (Dey et al. 2008; Prasad et al. 2003). The FEJ_1-4_ lines were derived from four ancestral populations (JB_1-4_), which served as controls in that experiment. Incidentally, one of these JB populations is the ancestor for the 32 populations used in the present study. For each FEJ*_i_* or JB*_i_* (*i* ɛ 1-4) population (represented by single vial cultures), there were four replicates each under HL and LH regimes. Thus, there were 16 FEJ populations and 16 JB populations that experienced the LH regime and similarly 16+16 that experienced the HL regime. The maintenance details of this 36-generation long experiment are given elsewhere (Dey et al. 2008) and are similar to the experimental protocol of the present study.

The rationale and the details of the model recalibration are presented in the SOM (Text S7). It should be noted here that the empirical *FI* values are those reported in Figure 2a of the earlier study (Dey et al. 2008) and are being re-plotted here only to facilitate comparison with the simulation results. The population size distribution data from these experiments is being reported for the first time here.

### 2.5 Simulating the dynamics of sex-biased populations

This part of the study consisted of three inter-related questions on the consequences of systematic departures from equal number of males and females in the population. We mention the simulations briefly here, with the details being relegated to Text S8.

a. Distortion of sex ratio and sex-specific culling: Here we simulated the behaviour of *FI* and average population size for two different values of fecundity values (high: *x_5_* = 85 and low: x*_5_* = 17) crossed with two values of *food* (high = 1.76 and low = 2.56). For sex ratio distortion, we varied the probability of an individual being female from 0.1 to 0.9, while for sex-specific culling, a fixed percentage (0% - 95%) of males or females were removed after the assignment of sex.
b. Sterile insect technique (SIT): In this method, a fixed number or proportion of sterile males are introduced into the population every generation. Following an established modelling strategy (Knipling 1959), we assumed the proportion of females producing eggs for the next generation to be a linear function of the fertile/sterile male ratio. Thus, proportion of egg laying females = min((*num_fertile*/(*β*×*num_sterile*)), 1), where *num_fertile* and *num_sterile* are the number of fertile and sterile males in the population, *β* is the competitive ability of the sterile males and min() is the minimum function. We studied the interaction of density-independent female fecundity with the amount of juvenile food available to determine the efficacy of SIT. The latter was estimated as the minimum number of sterile males required to ensure extinction of the population within an arbitrary number of generations (here 10) with a reasonably high success rate (here 90%). For this, we explored a large range of parameter values for density-independent fecundity (i.e. *adnut* × *x_5_*, range = 1 - 150, step size = 5) and larval food (*food*, range = 0.5 - 3.0, step size = 0.2). Note that strictly speaking, the product of *x_5_*, *adnut* and ln(*x_6_ + sen_adsize×m_c_*) should be called density-independent fecundity. However, as the values of *x_6_*, *sen_adsize* and *m_c_* are kept constant in these simulations, for the sake of simplicity, we continue to denote *adnut* × *x_5_* as density-independent fecundity. To avoid confounding extinctions due to intrinsic reasons with extinctions due to SIT, the efficacy of SIT was estimated only for those parameter combinations where the probability of an extinction event within first 10 generations in the absence of SIT is below 0.1 (Figure S10, see section 3.9 for discussion).

## 3. RESULTS AND DISCUSSION

### 3.1 Experiments: Larval and adult nutritional regimes interact to shape the adult dynamics

There was a significant interaction between the larval and adult nutrition (*F*_1, 28_= 17.92, *p*= 0.0002) suggesting that enhancing the fecundity of flies (through a supply of yeast) causes a much greater increase in *FI* when the amount of larval food is limiting (i.e. LL and LH) than when it not limiting (i.e. HL and HH) (Figure S1). This is because although both LL and LH experience substantial larval crowding, the greater fecundity of the LH flies leads to higher larval crowding even with moderate adult population sizes which, in turn, causes regular population crashes. On the other hand, even when there are population crashes, the greater fecundity of the LH flies ensures a high population size in the next generation. Together, these two effects ensure large amplitude oscillations in LH population sizes, and a substantially larger *FI* than the LLs (Tukey’s HSD *p* = 0.00016). This effect is also apparent from the alternation of negative and positive autocorrelations (Figure S5A), indicating population cycles with frequency of ∼ 2 generations. On the other hand, although the fecundity of females in the HH populations is larger than those in the HLs, the non-limiting amount of larval food ensures that the population crashes are only marginally more severe in the former. This leads to a much lower (although statistically significant; Tukey’s HSD *p* = 0.00017) increase in *FI* in the HH populations, compared to the HL populations (Figure S1).

### 3.2 Experiments: The differences in the dynamics of the populations are reflected in their population size distributions and *FI*

We began our investigation by examining the distribution of population size which is ultimately related to the temporal dynamics of populations (Figure S3). Both larval and adult nutritional levels were found to affect the population size distributions (Figure 1A, the white boxes). Specifically, when larval food is low, population size distributions have lower values of mean, median, 25^th^ percentiles and 75^th^ percentiles, compared to when larval food is high (*cf* LH with HH and LL with HL in Figure 1A). Interestingly, irrespective of the level of larval nutrition, providing yeast to the adults reduced the population sizes (*cf* LH with LL and HH with HL in Figure 1A). Moreover, in the LH and HH regimes (Figure 1A), the population size distributions are much more skewed to the right (i.e. median < mean), which is indicative of crashes in population numbers from various medium to high population sizes (see also figure 4 in Dey and Joshi 2013). All these observations are due to the fact that low levels of larval food or increased adult fecundity increase the larval crowding by reducing the per-capita food available to the larvae. Consequently, fewer larvae are able to acquire a body size greater than *m_c_*, which reduces the egg-to-adult survivorship, and hence the adult population sizes. Interestingly, the mean and the median population sizes were very close for the HL populations, but not so for the other three (Figure 1A). This showed that the population size distributions of HL had little or no skew, while the other three regimes exhibited positive skewness. This implied that in spite of having a larger average population size compared to the other three regimes, the HL populations exhibited lower amplitude fluctuations relative to their own mean population size. Thus, not surprisingly, the HL populations were found to have the lowest *FI* (Figure 1B) amongst the four regimes.

**Figure 1.**
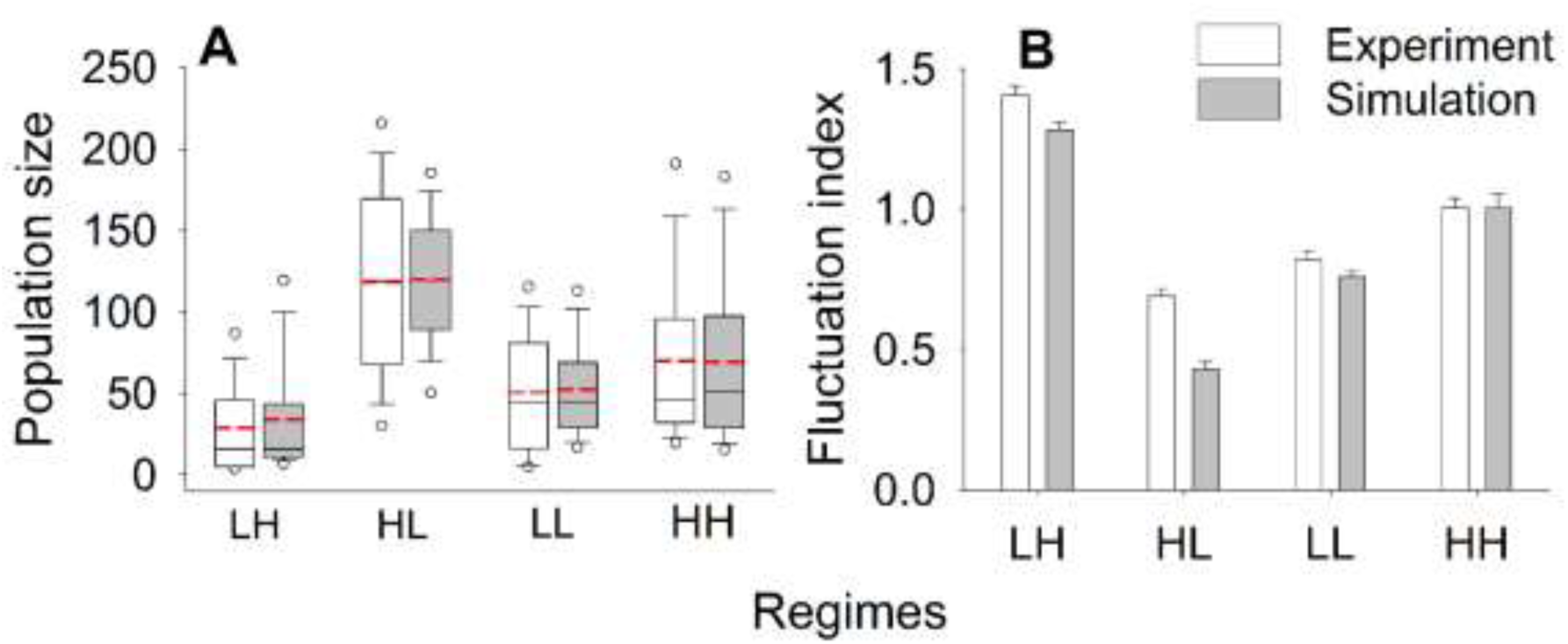
Population size distribution and constancy stability of experimental and simulated time-series. (**A**) Descriptive statistics of the population size distributions. Red dashed lines = means, thin black lines = medians, edges of the boxes=25^th^ and 75^th^ percentiles, whiskers=10^th^ and 90^th^ percentiles and the circles outside = 5^th^ and 95^th^ percentiles of the distributions. White boxes represent experimental data while grey shaded boxes are for simulated time-series. (**B**) Average (± SEM) *FI* of the experimental and simulated time-series in the four regimes. Both plots suggested a good agreement between the experiments and the simulations. The populations in the HL regime were the most stable with highest average population size while those in the LH regime were the least stable with lowest population size.

Post-hoc tests (Tukey’s HSD) on *FI*s in the four nutritional regimes revealed all pair-wise differences to be significant with the rank order: LH > HH > LL > HL (Figure 1B, the white bars). Although these four regimes have never been studied together to date, subsets of them have been studied in all kinds of combinations. Thus, it has been shown that in terms of constancy stability LH< HL∼HH (Mueller and Huynh 1994), LH< HL and LH< LL (Dey and Joshi 2013). Our results (Figure 1B) are in excellent agreement with all these studies except those of Mueller and Huynh (1994) who showed theoretically and empirically, that the constancy stability of HL and HH were not different. Moreover, Mueller and Huynh (Mueller and Huynh 1994) also predicted the average population size of the HH regime to be much larger than that of the HL regime, which also did not match our observations (Figure 1A). We resolve this issue later in this study (section 3.5) using our individual-based model of *Drosophila* dynamics.

### 3.3 Simulations: High level of correspondence between the experimental data and the model output

The simulation results (grey bars) matched the various salient features of population size distribution (Figure 1A) and population stability (Figure 1B) in the empirical observations in all four nutritional regimes. To the best of our knowledge, there are no models of *Drosophila* dynamics whose predictions have been verified in this detail with experimental data. This is more a reflection on the paucity of good quality long time series data, rather than any shortcoming on the part of the modelers. In fact, in the context of dynamics of laboratory populations of *D. melanogaster,* our 49 generation data-set is perhaps the second-longest in the literature in terms of number of generations.

Although the model did an excellent job in capturing the various aspects of the experimental data (*cf* Figure S3 and S4; also Figure S5), these details (and therefore the parameter values that lead to them) are obviously experiment-specific and shall vary across studies. Therefore, the usefulness of our model is more in terms of the mechanistic understanding that it generates about how the dynamics is affected by the interaction of various life-history and environmental variables. That was our next object of investigation.

### 3.4 Simulations: The effects of various life-history related traits on dynamics

#### 3.4.1 Hatchability (hatchability) and critical size (m_c_)

Our model predicted that with decreasing hatchability of the eggs up to moderate values, population *FI* decreases (i.e. constancy stability increases) in all four regimes (Figure 2A). This is because a reduced hatchability in generation *t* is conceptually equivalent to reduced fecundity in generation *t*-1, which is known to be a stabilizing factor (Mueller 1988). Moreover, reduced hatchability can actually improve average population size by reducing the extent of larval crowding (see Figure S7 for higher *hatchability* values). Thus, as expected, the destabilizing effects of increasing hatchability are more pronounced where the larval crowding is already very high (LH) and are mildest where larval crowding is the lowest (i.e. HL). Interestingly when hatchability values are very low (∼0.35 or less), further reductions lead to a decline in the average population size without greatly affecting the variation in population sizes over time (Figure S7). Since the expression for *FI* includes the average population size in the denominator, this leads to a minor increase in the *FI* values for very low values of hatchability.

**Figure 2.**
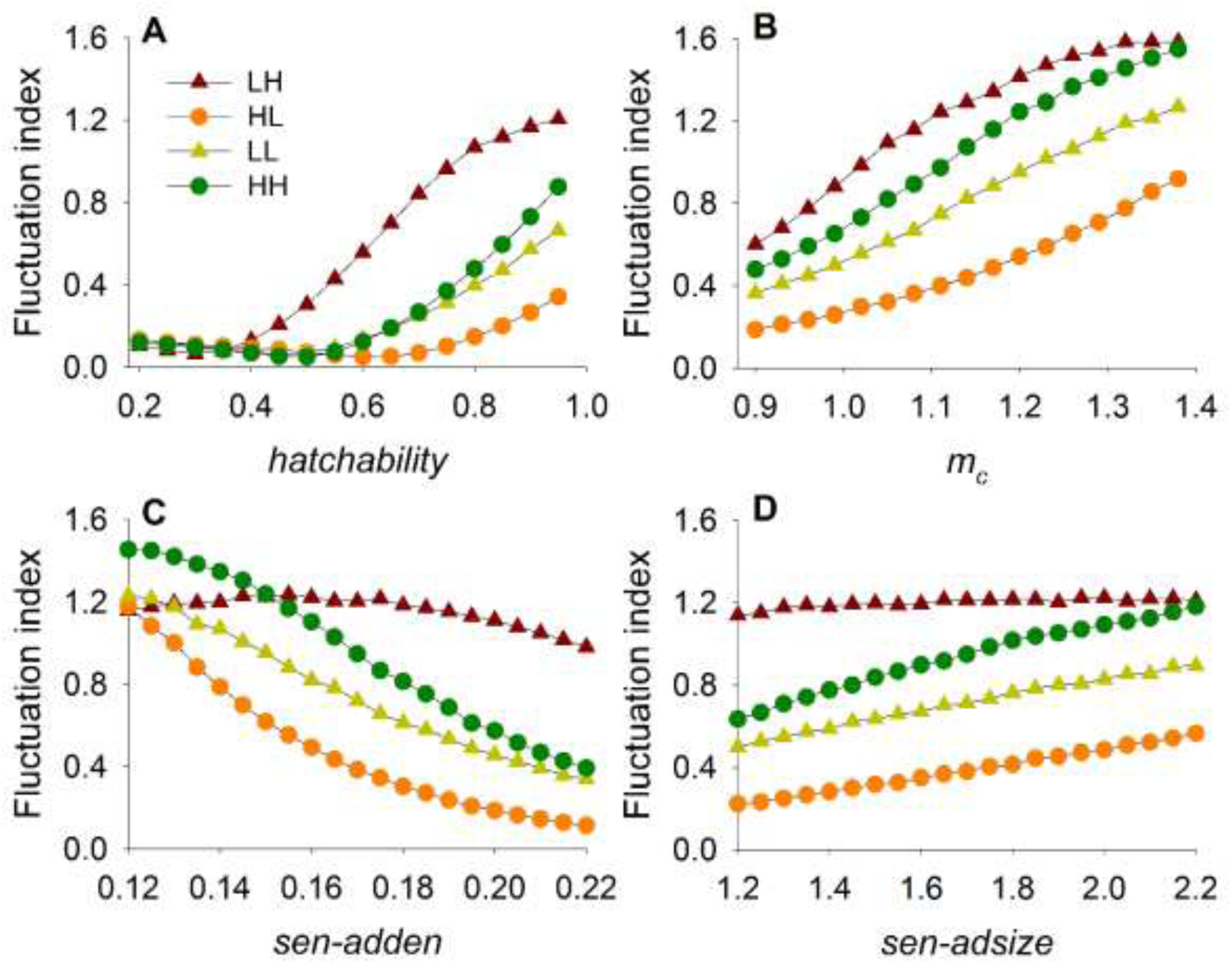
Effect of varying life-history related parameters of the model on constancy stability. Each point represents average (± SEM) fluctuation index of 100 replicates of 100-gen long simulated time series. Error bars are too small to be visible. (**A**) In all four regimes, as *hatchability* decreases, larval density also decreases and thus populations become more stable. (**B**) As critical size increases, the populations become less stable. (**C**) Increasing the sensitivity of female-fecundity to adult density (*sen_adden*) increases constancy stability in all regimes except LH. (**D**) Increasing the sensitivity of female-fecundity to adult body size (*sen_adsize*), decreases constancy stability in all regimes except LH. See section 3.4.2 for explanations for the anomalous behaviors in the LH regime.

Like hatchability, increasing larval critical size (*m_c_*) also decreases constancy stability (Figure 2B). This works in two ways. First, all else being equal, increasing *m_c_* means that fewer larvae would be able to attain *m_c_*, which would reduce larval survivorship. This is analogous to reducing survivorship through reduced larval food amount, which is a destabilizing factor. Secondly, increasing *m_c_* means that on an average, the surviving adults would be larger, which would translate into larger fecundity and thus, destabilize the dynamics. Conversely, decreasing *m_c_* is always expected to stabilize the dynamics (Mueller 1988), a prediction that we return to in section 3.6.

#### 3.4.2 Sensitivity of female fecundity to adult density (sen_adden) and adult body size (sen_adsize)

Increasing adult density is known to decrease female fecundity in *Drosophila melanogaster* (Mueller and Huynh 1994). In our model, *sen_adden* determines the strength of this effect, such that for same adult density, greater *sen_adden* results in lesser fecundity. This in turn enhances larval survivorship, by increasing the amount of food available per capita, which has a stabilizing effect on the dynamics. On the other hand, *sen_adsize* determines the strength of the positive correlation between body size and fecundity, such that increasing *sen_adsize* will increase fecundity, thereby reducing larval survivorship, ultimately leading to destabilized dynamics. In a nutshell, an increase in *sen_adden* and decrease in *sen_adsize* is expected to lead to a stabilization of the population dynamics. Our simulation results agreed with this prediction in all the regimes except LH (Figure 2C and 2D). In the LH regime, both *sen_adden* and *sen_adsize* seemed to have little effect on *FI*, even though reducing *sed-adden* and increasing *sen_adsize* caused the total egg number to go up (Figure S8A and S8C). The reason for this unintuitive behaviour was revealed when we investigated the effect of these two parameters on the egg-to-adult survivorship. Increasing *sen_adden* (Figure S8B), or decreasing *sen_adsize* (Figure S8D), hardly affected the egg-to-adult viability in the LH regime. This is because the reduced levels of larval food ensured that even with low fecundity, there was substantial larval crowding in this regime (as evidenced by the low levels of egg-to-adult viability) so that there was almost no effect of changing *sen_adden* or *sen_adsize* on larval mortality. As a result, the destabilizing effect of increasing fecundity was not seen in the LH populations. Another interesting observation from Fig 2C is that at low values (< 0.14) of *sen_adden*, HH has a much larger *FI* than LH, despite having similar levels of average egg number (Fig S8A). This is because for low values of *sen_adden*, the egg-to-adult viability of LH is lower than that of HH (Fig S8B), due to the fact that the amount of larval food for the former regime is much less than that of the latter. Consequently, greater number of adults can be supported by HH, which makes higher amplitude fluctuations in population sizes more likely, thereby increasing the values of *FI*.

The primary insight from the above observations is that even in the highly simplified dynamics under laboratory conditions, the environment can interact with the life-history related traits of the organisms to lead to counter-intuitive effects on the dynamics.

### 3.5 Simulations and Experiments: Population dynamics is shaped jointly by the quality and quantity of nutrition

As stated already, one of our empirical results did not match the observations of an earlier study (Mueller and Huynh 1994). We found that HL populations had greater constancy stability and average size than the HH populations whereas Mueller and Huynh (Mueller and Huynh 1994) reported that the HH populations had similar constancy stability but much greater average size than the HL populations. The primary difference between the two experiments was in terms of the amount of food given to the larvae. In the experiment of Mueller and Huynh (Mueller and Huynh 1994), the HL and HH larva got 40 mL of food in a 250 mL bottle while in our experiment the corresponding larva got ∼6 mL food in a 37 mL vial. Consequently, the adult population sizes in the HL and HH regime varied in the range of ∼40-240 in our experiment, but ∼ 400-1600 in the previous experiment.

To investigate whether the differences in the larval food amount could explain the observed discrepancies, we simulated the HH and HL regime for different levels of larval food, keeping all other parameters the same as in the earlier simulation. We found that as the level of larval food increased, the relationship between the population size distribution of HH and HL reversed (Figure 3A). Furthermore, with increasing value of larval food, the *FI* in the HH regime decreased and approached the value seen in the HL regime (Figure 3B). The underlying mechanisms behind these observations can be understood as follows.

**Figure 3.**
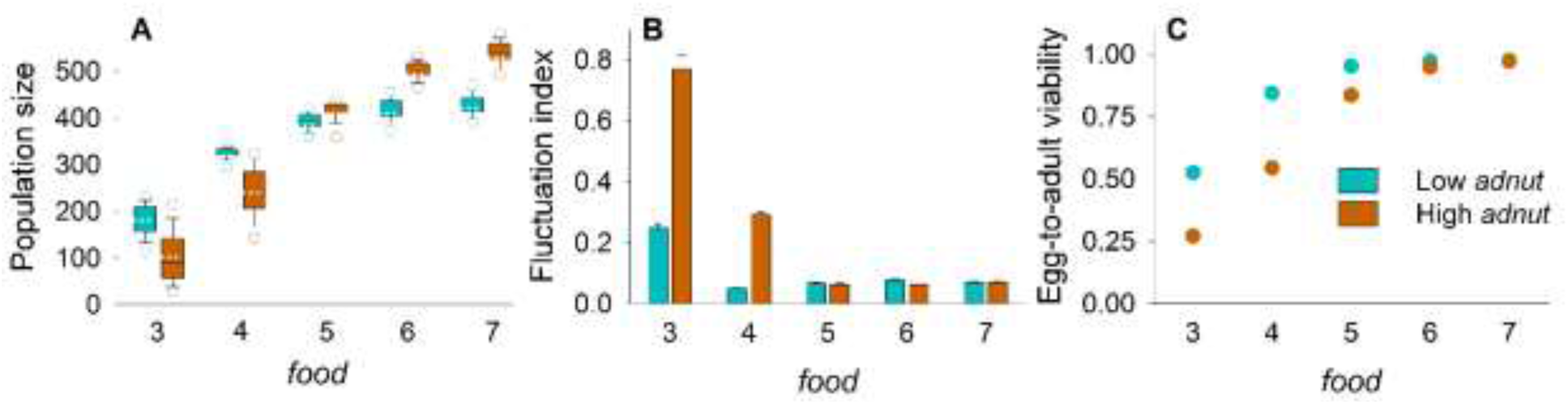
Simulations on effects of varying larval nutrition on population dynamics. (**A**) Population size distributions for the simulated time-series under low *adnut* (cyan) and high *adnut* (orange) conditions for different levels of larval food amount (*food*). White dotted lines = means, thin black lines = medians and the circles outside = 5^th^ and 95^th^ percentiles of the distributions. The relative positions of the population size distribution of low *adnut* and high *adnut* regimes reverses as the larval food amount increases. (**B**) Average (± SEM) fluctuation index of the low *adnut* and high *adnut* regimes become comparable, when the level of larval food is high. (**C**) Although the low *adnut* regimes have greater average (± SEM) egg-to-adult viability than the high *adnut* regime for low values of *food*, the viabilities become comparable as *food* increases. Error bars are too small to be visible here.

Due to the availability of yeast paste to the adults in the HH regime, the per-capita fecundity of the females is very high. Consequently, when the amount of larval food is relatively less (as in our experiment) there is larval crowding which reduces the survivorship in the HH regime. Therefore, with increasing levels of food, the survivorship increases (as in Figure 3C), which is manifested as increased population size in the HH regime (Figure 3A, orange boxes). The HL populations also face some amount of larval crowding at lower levels of food. However, since they do not have increased fecundity at high adult population sizes (due to the absence of yeast), the increase in population size plateaus off at a much lower level of food (Figure 3A, cyan boxes).

In order to visualize the effects of increased food amount on constancy stability, we need to appreciate that reduced larval crowding has two opposing effects on the dynamics. First, it stabilizes the dynamics by reducing larval mortality. At the same time, it can destabilize the dynamics by increasing the body size of the females at eclosion. As the larval food level increases, both these factors come into play. However, as there are upper bounds to both survivorship (=1) and the body size of the flies (= the physiological limit of body size), beyond a certain amount of larval food, both these factors cease to play a major role, and the *FI* in both regimes become similar. This can be clearly seen in Figure 3B and explains why in the presence of large amount of larval food, HL and HH populations have similar constancy, as reported previously (Mueller and Huynh 1994). When the amount of larval food is relatively small (while still being high, compared to LH or LL regimes), the destabilizing effect of reduced survivorship overpowers the stabilizing effect of diminished fecundity due to reduced body size. This is because there is a minimum value for the body size (= *m_c_*), which automatically places a lower bound on the fecundity of the flies irrespective of the level of larval crowding. Since the HH populations experience greater larval crowding than the HL populations, they are expected to exhibit lower constancy, as seen in our experiments (Figure 1B, white bars) and simulations (Figure 1B, grey bars).

In the *Drosophila* population dynamics literature, labels like LH and HL have typically been used as qualitative descriptors to signify the levels of larval crowding (highly crowded *versus* un-crowded) and state of adult nutrition (yeasted *versus* un-yeasted). As described in the Introduction section, this categorization has broad explanatory power in terms of the nature of the dynamics: LH leads to high amplitude oscillations while HL leads to relatively stable dynamics. However, the above comparison between the HL and HH regimes from the two different studies shows that changing just one environmental parameter (here, the absolute quantity of otherwise ‘high’ larval food) can lead to a rich array of dynamics. This again highlights how the actual values of the environmental parameters interact with life-history related traits in determining population dynamics.

### 3.6 Simulations and Experiments: Reduction in *m_c_* is one way for population stability to evolve

One of the predictions of our model is that decreasing *m_c_* should lead to stabilization of the dynamics (Figure 2B and Section 3.3.1). This prediction is consistent with earlier theoretical studies (Mueller 1988) and has been empirically validated using laboratory populations of *D. melanogaster* (Dey et al. 2008; Prasad et al. 2003). These earlier experiments used a population of flies (FEJs) that had evolved reduced *m_c_* as a correlated response to selection for faster development and early reproduction. Consequently, they were found to have reduced *FI* compared to the corresponding ancestral control populations (JBs). In order to see whether our model was capable of recovering the other features of the dynamics from the earlier experiment (Dey et al. 2008), we set a slightly lower value of *m_c_* for the FEJs and kept all other parameters the same as in the previous simulations (Table 1). Our model was again able to capture the trends in the distributional properties (Figure 4A) and the *FI* values (Figure 4B and 4C) of JB and FEJ populations in both regimes. The empirical data in Figure 1 and Figure 4 are from two completely independent experiments done at different times. The fact that our model is able to predict the major features of the latter data-set based on parameterizations done for a subset of the former shows that our parameterization was robust. However, we again emphasize here that the main focus of this part of the study was to gather insights about how the various life-history and environmental parameters interact, and the excellent quantitative match between the data and the model is essentially a secondary, though greatly encouraging, result.

**Figure 4.**
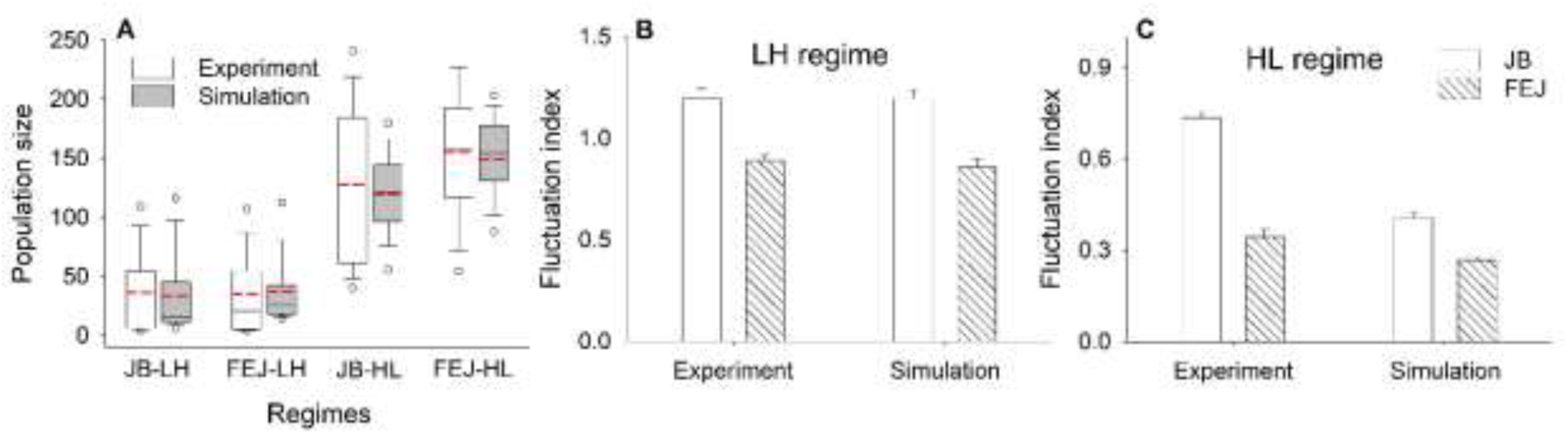
Validating model predictions on JB and FEJ populations. (**A**) Descriptive statistics of the population size distributions of experimental and simulated JB and FEJ populations. Red dashed lines = means, thin black lines = medians, edges of the boxes=25^th^ and 75^th^ percentiles, whiskers=10^th^ and 90^th^ percentiles and the circles outside = 5^th^ and 95^th^ percentiles of the distributions. White boxes represent experimental data while grey shaded boxes denote simulated time-series. Average (± SEM) *FI* of JB and FEJ populations corresponding to the experimental and simulated time-series under (**B**) LH and (**C**) HL regimes. Experimental data shows that in both the regimes, FEJs have lower *FI* than the JBs, as predicted by the model. Simulated FEJ populations capture well the empirical trends for population size distribution and constancy stability.

### 3.7 Comparison with experiments and models on other species

Since our model is based on fairly generic life-history features (see Text S3), it is reasonable to assume that it would be a good predictor of the dynamics across a wide range of taxa. For example, in another dipteran, the pitcher-plant mosquito (*Wyeomyia smithii*) increased larval food amounts reduces larval mortality and increases adult fecundity (Istock et al. 1975). Furthermore, in the same species, adult fecundity is positively and negatively affected by increased levels of adult food and adult crowding respectively (Istock et al. 1975). Since these are exactly the processes that govern the dynamics of *Drosophila* laboratory populations, it can be safely predicted that the dynamics of *W. smithii*, under different nutritional levels, will be well captured by our model. Unfortunately, we could not locate any studies on the population dynamics of this species to verify this prediction.

Paucity of studies in the literature was not the problem with the widely-studied crustacean genus *Daphnia*. In this system, increasing food concentration increases size at maturity and fecundity (Boersma and Vijverberg 1995; Rinke and Vijverberg 2005), as well as the slope of the body length-clutch size relationship (Rinke and Vijverberg 2005), which is equivalent to increasing the value of *sen_adsize* in our model. An IBM incorporating these relationships predicted increased amplitude of fluctuations in population sizes with increasing concentration of food (Vanoverbeke 2008), which agrees well with experimental observations (McCauley et al. 1999) and our results (Figure 2D). Simulation studies on another crustacean (*Armadillidium vulgare*) showed that increased level of food also increases the population size (Rushton and Hassall 1987), which is consistent with our observations (Figure 3A). It should be noted here, though, that all these *Daphnia* experiments and models deal with continuous culture systems, as opposed to the discrete generation systems investigated in our study. Moreover, in these systems, the adults and the juveniles share the same food, whereas in our model, the effects of the larval food (determined by the parameter *food*) are different from the effects of the adult nutrition (modeled using *adnut*). In spite of these non-trivial differences, the primary mechanism by which food amounts affect the life history, and thereby the dynamics, remain alike.

Moving away from the arthropods, not surprisingly, leads to more serious departures from the insights gathered from our study, although some broad patterns are still discernible. For example, an IBM for the fish yellow perch (*Perca flavescens*) showed that increased food (forage fish) amount leads to larger body size, higher fecundity and lower adult abundance (Rose et al. 1999). However, in the same study, when the amount of food was increased for another fish, walleye (*Sander vitreus*), the model predicted larger body size, lower fecundity and higher adult abundance. This difference was partly due to the fact that the life-history parameters and the ecological food-bases of the two fishes were very different. Although the authors did not report the effects of these food manipulations on the population dynamics, it is clear that changing the levels of food can affect the population size, which appears to be a fairly robust observation in this context. In a comprehensive review covering 138 species, including reptiles, birds and small mammals, Boutin reported that adding food to the environment typically leads to 2-3 fold increase in population density but little or no change in the population dynamics (Boutin 1990; although see Klenner and Krebs 1991 and references therein). More importantly, litter or clutch sizes are hardly affected by food supply, except when there is severe food shortage due to natural reasons or severe overpopulation (Boutin 1990).

There can be several reasons for the differences between the observations of these vertebrate studies and ours. First, there is no reason to expect that a model based on arthropod life history can accurately capture the population dynamics of vertebrates. Second, it is very difficult to estimate the parameter space for the vertebrate systems that is analogous to the parameter space used in our model. Third, all the vertebrate studies quoted here are on natural populations with multiple sources of food and other ecological interactions, at least some of which can interact with the effects of food manipulation in non-intuitive ways (e.g. Klenner and Krebs 1991).

The above account suggests that there are substantial differences between how food levels affect the dynamics of insects / crustaceans on one hand and higher vertebrates on the other. Not surprisingly, our model is a better descriptor of the former. After establishing what our model could and could not do, we used it to investigate the interaction of resource availability with life-history traits and demographic factors in shaping dynamics and stability of stage-structured populations.

### 3.8 Simulations: The effects of sex-ratio and sex-specific culling are greatly influenced by fecundity but not by levels of juvenile food

Theoretical studies have typically neglected the effects of sex ratio on the stability of population dynamics (although see Johnson 1994), an important lacuna that we address here. We found that when the baseline fecundity is high, the *FI* of the population is highest when the sex ratio is close to 1:1 and decreases when the sex ratio becomes more skewed (closed circles in Figure 5A). This is consistent with the observation that population growth rate is typically largest either when both sexes are present in equal proportions or when the proportion of females is slightly higher (Jenouvrier et al. 2010; Miller and Inouye 2013). However, although equal or female-biased sex ratios are also expected to maximize the average population size (Rankin and Kokko 2007; Schmickl and Karsai 2010), our model showed that the maximum population size was reached when the proportion of females in the population was ∼15%, i.e. when the population was male-biased (closed circles in Figure 5B). This general pattern was not altered irrespective of the amount of food given to the larvae (*cf* Figure 5 with Figure S9).

**Figure 5.**
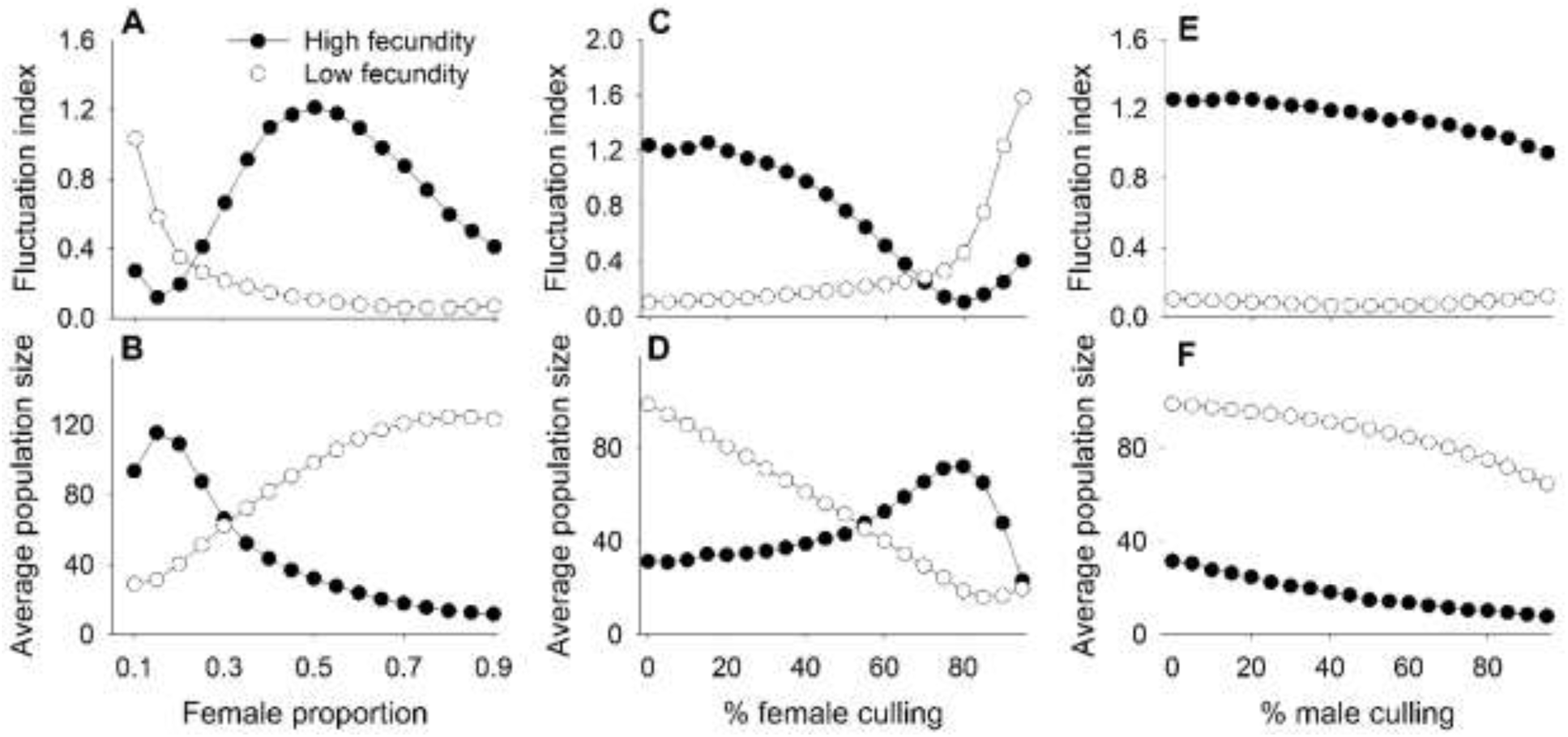
Effect of varying sex-ratio and sex-specific culling on population dynamics for low larval nutrient scenario. **A)** Fluctuation index and **B)** Average population size as a function of the expected fraction of females in the population. **C)** Fluctuation index and **D)** Average population size as a function of the percentage of females culled. **E)** Fluctuation index and **F)** Average population size as a function of the percentage of males culled. Closed and open circles denote high and low levels of density-independent adult fecundity. See section 3.8 for explanations. Each point represents average (± SEM) fluctuation index or population size of 100 replicates of 100-gen long simulated time series. Error bars are too small to be visible.

To explain this observed discrepancy, we note that many of the previous studies on effects of sex ratio assume the mating success of the females to be a function of the number of available males (Jenouvrier et al. 2010; Rankin and Kokko 2007; Schmickl and Karsai 2010). Although applicable for strictly monogamous species, this assumption need not always be true for insects like *D. melanogaster* where both males and females mate multiple times (although see Vahl et al. 2013), and a single mating can sustain high lifetime fecundity of a female, unless the proportion of males is extremely low. That is the reason our model assumed that all females contributed to the egg numbers of the next generation, irrespective of the number of males in the population (we relax this assumption in the next section). Consequently, male numbers can only have negative effects on the number of eggs in the next generation through the density-dependent reduction in fecundity. On the other hand, each female in the population has a positive effect on the number of eggs in the next generation through its fecundity, and a negative effect through its contribution to the density-dependent reduction in fecundity (eq. 4). The sex ratio leading to the maximum population size is thus determined as an interaction between these two opposing effects. When female fecundity is high and not male-limited, this number is much lower than what is expected when these two conditions are not met. This explains why, in our model, the maximum population size is attained under male-biased conditions. When the number of females in the population is increased beyond this threshold, the number of eggs in the next generation increases, which, in turn, results in more intense larval crowding. Conceptually, this is similar to boosting the fecundity by supplying yeast to the adults which reduces the average population size (Figure 1A), as can be seen by comparing LL with LH and HL with HH in Figure 1A.

When the density-independent fecundity was low (open circles in Figure 5A and 5B) there was monotonic increase and decrease in average population size (Figure 5B) and *FI* (Figure 5A), respectively, with increasing proportion of females in the population. This trend was clearly different from the high fecundity case (*cf* open circles and closed circles in Figure 5A and Figure 5B), indicating that fecundity itself can alter the effects of sex ratio on population dynamics. To the best of our knowledge, this effect has previously not been reported in the literature and can be understood as follows. When the per capita female fecundity is low, the number of eggs in the next generation is reduced and the average size of the population is therefore limited more by the total number of eggs produced than the larval density-dependent mortality. Moreover, under such circumstances, the populations are more vulnerable to demographic stochasticity, which, in conjunction with low average population size, leads to an increase in *FI*. Consequently, with increases in proportion of females, both the average size of the population and the population stability increases.

#### Sex-specific mortality or culling of the adult individuals

In a population, sexes often suffer unequal mortality rates. This can happen due to, *inter alia*, sex-biased predation (Boukal et al. 2008) or human preference for a given sex for commercial or other purposes. We modeled this phenomenon using our model by explicitly culling a fixed percentage of a sex in each generation and, not surprisingly, found that female (Figure 5C-D) and male (Figure 5E-F) culling have very different effects on the dynamics. The patterns observed in Figure 5C-F can be explained by noting that female culling increases the ratio of males in the population and vice versa, and the subsequent effects on *FI* and average population size can be deduced based on Figure 5A-B. Incidentally, in case of male culling, the density-dependent reduction in fecundity also decreases due to reduced population size. As a result, the drop in FI is much slower for high fecundity in Figure 5E than it is expected from Figure 5A alone. Thus, the reduction in average population size in Figure 5F is a trivial consequence of male culling. However, it is interesting to note that when the female fecundity is high, culling of females can increase the average population size, a phenomenon that is termed as the Hydra effect in the population dynamics literature (Abrams 2009). However, when female fecundity is low, increasing the fraction of females culled reduces population size, which is consistent with recent theoretical and empirical data on the beetle *Callosobruchus maculatus* (Snyder et al. 2014). Moreover, although changing the amount of juvenile food led to changes in the numerical values of the average population size and *FI*, the corresponding patterns remained unaltered (*cf* Figure 5C-F and Figure S9C-F), suggesting that the effects of changes in sex-ratio on population dynamics are not altered by juvenile nutrition.

To summarize, different amounts of juvenile food affects the interaction of sex ratio and population dynamics quantitatively but not qualitatively. However, in species like *Drosophila*, where adult fecundity can be increased by giving external food (equivalent to increasing *adnut*), adult food levels will play a role in determining the qualitative and quantitative effects of unequal numbers of sexes on the dynamics.

### 3.9 Simulations: Fecundity and nutrition levels interact to determine the efficacy of Sterile Insect Technique (SIT)

One of the major applications of skewed sex-ratios has been in the context of SIT, wherein a large number of sterile individuals (typically males) are released into an insect pest population (Klassen 2005). These sterile individuals compete with the fertile individuals, thus reducing their reproductive fitness, which ultimately leads to effective suppression of population density or complete eradication of the pests (Knipling 1955). This method has been successfully used to eradicate several pest species like melon fly, Tsetse fly, screwworm, Mexican fruit fly and West Indian fruit fly (see Dyck et al. 2005 for a comprehensive review). Not surprisingly, the consequences of releasing large number of sterile males have been extensively investigated in the theoretical population dynamics literature (reviewed in Barclay 1980) primarily using simple non-linear models (e.g. Berryman et al. 1973). These studies have typically not incorporated the details of life-history or environmental variables like food amount on the efficacy of SIT. In fact, we could locate very few individual or agent-based models on SIT in the literature (Isidoro et al. 2009; Lin et al. 2015; Stone 2013). Simple models predict that when the fecundity of females is high, introducing sterile males can increase the pest population size (Berryman et al. 1973), whereas the present study shows that changes in larval food amount can affect the dynamics of populations. Therefore, we used our model to explore the interaction of the amount of food and fecundity of the pests on the expected number of individuals needed for a successful pest eradication using SIT.

Following prior studies (Knipling 1959), we assumed that the number of females that get fertilized and lay eggs is a function of the ratio of fertile to sterile males in the population (see section 2.5). As noted earlier, neither our fly populations nor our model-derived time series underwent too many extinctions. However, for this set of simulations, we needed to consider parameter values of food and density-independent fecundity (i.e. *adnut* × *x_5_* in equation 3) that were very different (greater as well as lesser) than what was applicable for our fly populations. Consequently, the extinction rates no longer remained negligible which in turn meant that sterile male induced extinctions needed to be distinguished from endogenous extinctions. Therefore, we first simulated the probability of extinction over 10 generations for various combinations of fecundity and food values (Figure S10). From this set, we used only those fecundity-food combinations which had less than 10% probability of going extinct without any perturbation to estimate the minimum number of sterile insects to be introduced that could lead to population extinction within 10 generations with a 90% probability (Figure 6). Thus, the lower the number of males needed to induce extinction, the higher was the efficiency of SIT. We found that when density-independent fecundity is <∼40, changing the amount of food had no effects on SIT efficiency. However, beyond this range, the efficiency of SIT reduced with the amount of food available, and the effect became more prominent as fecundity increased. The reason for this observation can be understood when we consider the effects of increasing the amounts of food for different fecundity levels. When fecundity is low, larval crowding is less and therefore, increasing the amount of food does not increase the population size. Consequently, the number of sterile males needed to bring extinction is also unaffected. However, when the fecundity is high, there is considerable larval crowding. Consequently, increasing the amount of food increases the average population size (as seen in Figure 3A) which in turn reduces the efficiency of SIT. Thus we find that the efficiency of SIT depends on an interaction between the amount of food available to the larva and its density independent fecundity (which can be altered by environmental factors). To the best of our knowledge, this interaction has not hitherto been reported. It should be noted here, that for our flies, the values of density independent fecundity (i.e. *adnut* × *x_5_*) considered were 85 and ∼125 for un-yeasted and yeasted conditions respectively. Therefore, the observed interaction between food and fecundity happens in a parameter range for fecundity (>∼40) which is very relevant biologically. Another point regarding these results is the order of magnitude of the population sizes. In nature, pest population sizes would typically be in orders of 10^5^-10^7^ whereas in our simulations, the population sizes were in orders of 10^1^-10^2^. This is because it is known that SIT is typically much more effective at low population sizes and therefore any application of SIT in the field is typically preceded by a chemical treatment that greatly reduces the population size (Klassen 2005). Thus, although the population sizes in our simulations are smaller than what would be experienced in a real-world application of SIT, they are off by no more than 1-2 orders of magnitude. Nevertheless, we believe that our results are still relevant to SIT application since the purpose of our study was not to model the application of SIT to a particular real world population of a specific insect pest species, but rather to derive insights about how outcomes of sterile male release could possibly be affected by nutrition × fecundity interactions.

**Figure 6.**
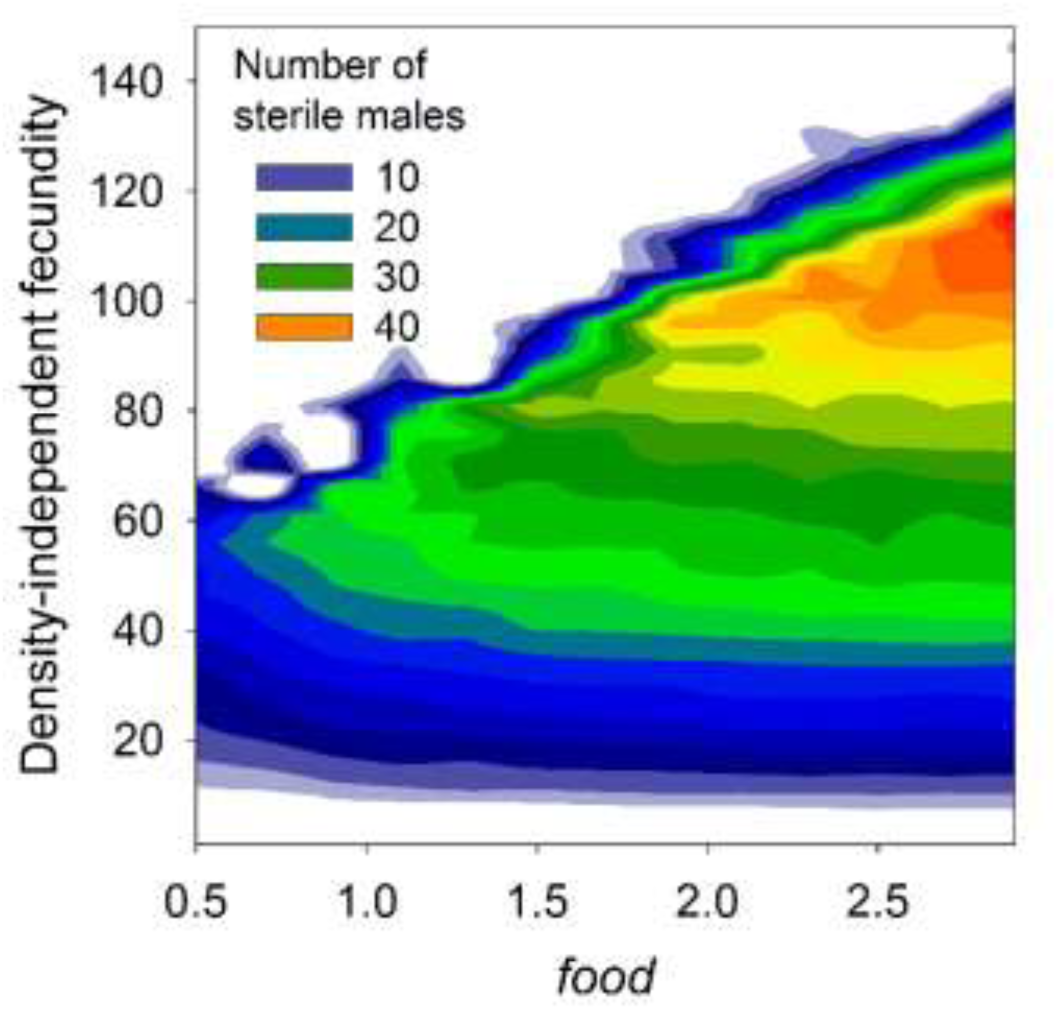
Effect of varying food and density-independent fecundity of the model on sterile male technique. For each combination of *food* and density-independent fecundity, this graph denotes the minimum number of sterile males needed to induce extinction within 10 generations with 90% certainty. Each point here is an average value obtained from 100 replicate simulations and plotted according to the adjacent colour index. Broadly, when fecundity is low, required number of sterile males does not depend on the amount of available food. But, for higher fecundity values, food and fecundity together determine the required number of sterile males needed for the desired level of extinction. For most levels of food studied, the required level of extinction was hardest to achieve (i.e. maximum number of sterile males were needed) at intermediate levels of density-independent female fecundity. For this analysis, we have considered only those food × fecundity combinations, where the basal probability of extinction is <10%.

## 4. CONCLUSION

Mathematical modeling of the dynamics of laboratory populations has a long and venerable history (Kingsland 1995; Mueller and Joshi 2000) and has been extensively done for several model systems like *Tribolium* (Costantino et al. 1997), *Callosobruchus* (Tuda and Shimada 2005), protists (Holyoak et al. 2000), mites (Benton and Beckerman 2005) etc. Depending on the objectives of their investigation, these studies have employed different kinds of modeling tools, ranging from simple deterministic difference equations, to coupled differential equations and individual-based models (reviewed in Mueller and Joshi 2000). The value of our study is first in the close correspondence between empirical observations and simulation results, and second in terms of the insights gained regarding the interaction of the environmental factors (larval and adult food level) with life-history related traits to determine population dynamics and stability. Our model was able to replicate several predictions about the dynamics of insects and crustaceans. Given that these two taxa represent about 70% of all animal species on earth (Zhang 2011), one can be reasonably confident that the general insights derived from this study are broadly applicable.

## ACKNOWLEDGEMENTS

We thank N. G. Prasad, Mallikarjun Shakarad and M. Rajanna for help in conducting the experiments, and Neelesh Dahanukar for helpful discussion regarding data analysis. For financial support, ST thanks JNCASR, Bengaluru, for a Project Oriented Biological Education (POBE) fellowship, and the Council for Scientific and Industrial Research, Government of India, for a Senior Research Fellowship. AJ thanks the Science and Engineering Research Board of the Department of Science and Technology, Government of India, for support through a J. C. Bose National Fellowship. This study was partly sponsored by an extra-mural grant from the Council of Scientific and Industrial Research, Government of India, and in-house funding from IISER-Pune and JNCASR, Bengaluru.

## Supplementary online material

### Text S1. Laboratory ecology of *Drosophila melanogaster*

In laboratory cultures of *D. melanogaster*, if the larval crowding is high, the mean amount of food available per larva is reduced. As a result, a large proportion of larvae are unable to attain the critical body mass needed for successful pupation, thus increasing larval mortality (Bakker 1961). Since the body size of the adults depends mainly on the amount of resources gathered during the larval stage, the adults emerging out of crowded cultures are generally small in size (Marks 1982) and exhibit low fecundity (Chiang and Hodson 1950). Adult fecundity is also reduced with increasing density of adults in a culture and this is generally attributed to increased interference with egg laying (Pearl 1932). Interestingly, this negative effect of adult density on fecundity can be ameliorated to a substantial degree by supplying the adults with excess amount of live yeast paste (Mueller and Huynh 1994). Since survival and fecundity are the major factors affecting the growth rate of a population, it seems plausible that these three density-dependent feedback loops ― effects of larval crowding on larval survivorship and subsequent adult fecundity, and effects of adult crowding on adult fecundity ― play a major role in determining the dynamics and stability of *D. melanogaster* populations in the laboratory (Mueller and Joshi 2000).

### Text S2. Experiment

The experiment was comprised of thirty-two populations of *D. melanogaster*, each represented by a single vial (9 cm h × 2.4 cm dia.) culture. These populations were derived from a long standing, large outbred population (JB_1_), maintained on a 21-day discrete generation cycle. Details of the ancestry and maintenance protocol of the JB populations can be found elsewhere (Sheeba et al. 1998), and are not germane to this study. These 32 populations were randomly allotted to one of four nutritional regimes, such that there were eight populations per regime. Following established norms (Mueller and Huynh 1994; Mueller et al. 2000) these regimes were called HH, HL, LH, LL — where the first letter indicates the quantity of larval food and the second letter represents the status of adult nutrition. In case of larval food, H and L denoted ∼6 mL and ∼2 mL of banana-jaggery medium per vial, respectively, whereas in the case of adult nutrition, H and L referred, respectively, to the presence and absence of live yeast paste supplement to banana-jaggery medium. Thus, for example, HL denotes a nutritional regime comprising of ∼6 mL medium per vial for the larvae, but no live yeast paste supplement for the adults, and so on.

Each population was initiated (generation 0) with eight male and eight female flies, and from this point onwards (except for extinction) there was no direct control on the number of adults in a vial. After oviposition in the vial for 24 hours (day 0), the adults were counted and discarded and the eggs formed the next generation. Once the adults started eclosing in these vials, they were transferred to adult collection vials every day with a change of medium every alternate day. Strict vial-to-vial correspondence was maintained between the egg vials and their corresponding adult collection vials. The process of adult collection continued until 18 days after day 0, after which the flies were conditioned for three days in the presence / absence of live yeast paste. The live yeast paste is known to boost the fecundity of the females (Chippindale et al. 1993) and reduce the effect of adult density on adult fecundity (Mueller and Huynh 1994). On day 21 after egg collection, the adults were transferred to fresh food vials containing ∼2 mL or ∼6 mL of banana-jaggery medium for a duration of 24 hours. After this, the adults were counted and discarded, while the eggs laid during this period formed the next generation. If there were no adults in a population, then an extinction event was recorded and the population was rescued by allowing four female flies from the ancestral JB_1_ population to lay eggs for 24 hours. All extinction were recorded in the adult time series as 4 individuals (i.e. the number of rescuing females).

The complete details of this experiment have been reported in the PhD thesis of one of the authors (Dey 2007).

### Text S3. Model formulation

The model can be divided into two modules: pre-adult and adult. For a given generation *t*, the pre-adult module takes the number of eggs and the total amount of larval food as input and computes the number of viable adults and the body size of each of those adults as an output. The output of the pre-adult module and the nature of the adult food available act as inputs for the adult module and the output is the total number of eggs produced that form the input for the pre-adult module in generation *t +1*. Thus, although our model produces the adult numbers in each generation, structurally it is an egg-to-egg recursion. This modelling strategy has been employed earlier (Mueller 1988), and is preferred over an adult-to-adult recursion. This is because, due to density-dependent mortality, for a given amount of larval food, the relationship between adult numbers and the corresponding number of eggs from which they have arisen is single-humped (Chiang and Hodson 1950). Consequently a given number of adults can arise from very different number of eggs (Prout and McChesney 1985). Thus, for example, assuming say 10% mortality at low crowding, 10 eggs will always lead to ∼9 adults. However, if one sees 9 adults, this could have arisen from 10 eggs (assuming 10% mortality at low crowding) or 100 eggs (assuming say 91% mortality at high crowding). Thus, tracking the adult numbers is never sufficient for the purpose of modelling *Drosophila* dynamics (Prout and McChesney 1985).

#### S3.1 Pre-adult module

##### S3.1.1 Step 1: Obtaining number of larva (numlarva) from the number of eggs (numegg)

This module starts with a given number of eggs (`numegg`) and assumes that only a fixed fraction (*hatchability*) of them will hatch into larvae, due to density-independent mortality. Thus, the number of viable larvae is given by

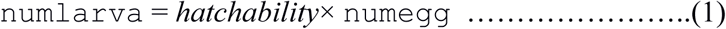

where 0 ≤ *hatchability* ≤1. Reasons like intrinsic poor viability of the eggs, environmental stresses (say some toxins in the environment) or other ecological/random stochastic factors, can make *hatchability* < 1. However, under normal laboratory conditions, hatchability generally remains above 0.9 (Bakker 1961). Therefore, we have taken *hatchability* = 0.98 and kept it same for all the simulations throughout the study (unless explicitly mentioned otherwise). Note that here we consider *hatchability* as a density-independent parameter because there is no experimental evidence to suggest that hatchability is affected by egg density. However, *hatchability* can be easily made a function of egg or adult density, without affecting the other parts of the model.

##### S3.1.2 Step 2: Obtaining the body size of each larva

In a *Drosophila* culture, the newly hatched larvae eat the larval food provided and grow in size and body mass. Although in the strict sense, size and mass are different quantities, for the sake of simplicity, we use them interchangeably to indicate biological growth. Due to among-individual variation in traits like larval feeding rate, food-to-biomass conversion efficiency etc., a distribution of larval body sizes ensues at the end of the larval growth period (Bakker 1961). When the number of larvae in the food is increased, the amount of food available per larva, on an average, is reduced which, in turn, reduces the average body-size attained at the end of the larval stage (Chiang and Hodson 1950; Miller and Thomas 1958). Following a previous study (Bakker 1961) we assumed the distribution of larval body sizes at the end of feeding to be normal with a mean (mean_size) that was an increasing function of the total amount of larval food (*food*), but a decreasing function of the number of larvae (`numlarva`). Specifically,

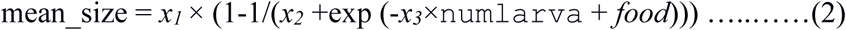

where *x_1_*, *x_2_* and *x_3_* are parameters (non-negative constants) and exp is the exponential function. The function is a logistic function of ‘*food - x_3_* × `numlarva`’, which has an upper limit *x_1_* and lower limit *x1*(1-1/*x_2_*). Thus, theoretically, *x_1_* could be defined as the maximum possible average body size (theoretically attained when number of larvae is very small and amount of food is close to infinite) and 1/*x_2_* as the maximum fractional reduction in average body size (theoretically attained when the number of larvae is infinitely large and the amount of food is vanishingly small). We arbitrarily assigned *x_1_* = 2.5 and *x_2_* = 1. The parameter x_3_ determines the rate of decline of mean_size with increasing the number of larvae for a given value of *food*. The values of *x_3_* and low-high values of *food* were calibrated systematically (see Text S4) to match the observed empirical pattern of larval viability and body size for different egg densities with a fixed amount of larval food (Chiang and Hodson 1950; Rodriguez 1989). Note that here our intention was to adopt a function that can mimic the general qualitative features of the relationship between mean_size*, food* and `numlarva`. Equation 2 is just one function that fulfills this criterion and potentially many others would have also served our purpose (for example, logistic function of *food/*`numlarva` or variants thereof).

For the sake of simplicity, standard deviation (*sigma_size*) of the body-size distribution was kept as a density-independent constant (=0.45). Computationally, once `numlarva` is calculated from equation 1, each larva is assigned a body size value by drawing random numbers from a normal N(mean_size, *sigma_size^2^)* distribution. The absolute function (abs(*x*) = -*x* if *x* < 0 else *x*) is used to avoid any negative value of body size.

##### S3.1.3 Step 3: Critical mass cut off and adult body size

In order to complete metamorphosis and become an adult, *Drosophila* larvae, like those of many other holometabolous insect species (Davidowitz et al. 2003), need to attain a critical minimum size before pupation (Bakker 1961; Robertson 1963). To incorporate this phenomenon into our model, we considered a (deterministic) cut off, critical size (*m_c_*), to be a density-independent constant (following Mueller 1988) and compared the size of each larvae against it. All larvae whose body sizes were less than *m_c_* were considered to have failed in becoming adults. The number of remaining larvae was considered to be the adult population size (`numadult`) of the current generation.

Empirical studies indicate a positive correlation between larval and adult body size in *Drosophila* (Bakker 1961). Therefore, we considered adult body size to be a linear function of larval body size, i.e.

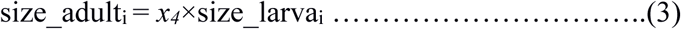

where size_adult_i_ and size_larva_i_ denote the body size of the *i*^th^ larva and the corresponding adult respectively (size_larva_i_ > *m_c_*) and *x_4_* is a parameter (= 1.0 for the sake of simplicity), which maps larval size to adult size. Note that *x_4_* is necessarily a positive quantity as it relates the body size of larvae to those of adults (both of which are positive). The nature of equation 3 ensures that heavier larvae lead to heavier adults, irrespective of the value of *x_4_*.

Thus, the pre-adult module takes a life-history variable (`numegg`) as an input and gives two life-history related variables, number of adults (`numadult`) and the distribution of the adult body sizes (size_adult_i_), as output.

Recently, it has been discovered that *Drosophila* larvae can exhibit cannibalism under conditions of extreme food deprivation (Vijendravarma et al. 2013). However, we chose not to incorporate this phenomenon in the model since the extent of cannibalism among the larvae under the level of crowding found in our populations is still not known. More critically, there is no evidence till date to indicate that this is a density-dependent phenomenon. If we assume larval cannibalism to be density-independent, then this phenomenon can be easily incorporated into our model by multiplying `numadult` with another constant.

#### S3.2 Adult module

##### S3.2.1 Step 1: Assigning a gender to each individual

The first task in this module is to assign a gender to each adult individual. In this study (unless explicitly mentioned otherwise), sex-ratio was considered to be independent of adult numbers and was always taken to be 1:1, so each adult was assigned to be female with probability 0.5 and male otherwise. However, due to the inherent stochasticity of the process, the realized sex ratio could deviate from 1:1, which is biologically realistic, particularly in small populations of the kind that we are studying. *Drosophila* is a sexually dimorphic species with the females being significantly larger than the males. Therefore, ideally, only the heaviest individuals should have been designated as females. However, we ignore this complication in our model and assign sex irrespective of adult body size.

##### S3.2.2 Step 2: Calculating the total number of eggs produced by all females

After the assignment of sex, fecundity of the females is computed based on their body size and current adult density. In many holometabolous insects, including *Drosophila*, fecundity or egg laying ability is positively correlated with the body size of the females (Honěk 1993). Prior studies have assumed that the fecundity of *Drosophila* females scale linearly on a ln-ln scale (Mueller 1988), which implies that slight increases in body size values in the upper range can lead to large increases in fecundity. This is somewhat unrealistic as larger flies would also require to expend substantial amount of energy in body maintenance and therefore it is more likely that the rate of increase of fecundity would eventually slow down with increasing body size. This relationship could be modelled in many ways and we chose to use the logarithmic function to represent the effects of female body size on adult density-independent fecundity. Finally, live yeast paste is known to boost female fecundity irrespective of the density (Mueller and Huynh 1994). This phenomenon, is incorporated by adding a density-independent constant (*adnut*) which denotes the fecundity boost due to yeast supplement to the adults (for not yeasted, *adnut* = 1; for yeasted, *adnut* >1). Taken together, the adult density-independent component of fecundity of the *i*^th^ female can be represented as:

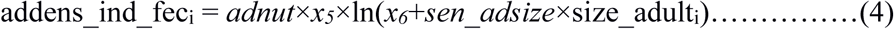

where *sen_adsize* is the strength of relationship between female-fecundity and adult body size and *x_5_* and *x_6_* are scaling parameters. It should be noted here that in the above formulation, two of the constants (*adnut* and *x_5_*) can easily be combined to create a single constant. However, we refrain from that in order to retain the ease of biological interpretation.

Another important factor that reduces per capita female fecundity in insects is adult density (Mueller 1988; Rich 1956). Following an earlier study (Mueller 1988) we modelled this relationship as:

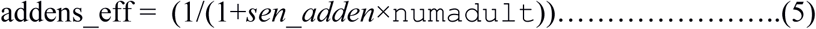

where *sen_adden* is the sensitivity of female-fecundity to adult density. This hyperbolic function denotes negative feedback of adult density on adult fecundity, a relationship which is found not only in *Drosophila* (Mueller and Huynh 1994; Rodriguez 1989) but in many other species as well (Băncilă et al. 2016; Michael and Bundy 1989). This negative feedback automatically makes population growth negatively density dependent and such negative feedback is known to be a necessary condition for population regulation (Turchin 1999). The values of *x_5_*, *x_6_*, *sen_adsize* and *sen_adden* were determined by an extensive search over the parameter space in order to match the experimentally observed fecundity of a healthy female fly as well as to satisfy a previously observed pattern of female fecundity versus adult density for both yeasted and un-yeasted conditions (Fig 2 in Mueller and Huynh 1994, Fig 6.5 in Mueller and Joshi 2000).

Combining equations 4 and 5, the fecundity of the *i*^th^female is given as:

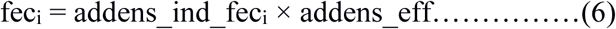

such that the number of eggs in the next generation,

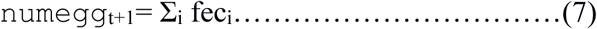

Thus, the adult module takes two life-history related parameters from the output of the pre-adult module and returns the number of eggs in the next generation as the output. This output then serves as the input for the pre-adult module for the next generation, and thus the iterations continue.

An earlier version of this model, and its correspondence with the experimental data, has been reported in the Master’s Thesis of one of the authors (Tung 2012).

### Text S4. Model calibration

The calibration of the individual-based model was attained in three sequential steps.

#### S4.1 Step 1. Calibrating the pre-adult module

Previous studies have shown that the value of *m_c_* is around half the maximum larval size that is physiologically attainable (reviewed in Prasad and Joshi 2003). Since we had already fixed the maximum value of the mean larval size (i.e. *mean_size*) at 2.5, we started with an initial value of *m_c_* = 1.25. Further, we arbitrarily started with *x_3_* = 0.01. We then simulated steps 1 and 2 combined (see Text S3.1) to obtain the effects of `numegg` (range 1-1500) on larval viability and larval body size, for different amounts of *food* (range 0.5-6.0). The upper limits on the ranges of `numegg` and *food* roughly correspond respectively to the upper ranges in egg numbers typically seen and the amount of food (in mL) typically used in *Drosophila* vial populations in the laboratory (Dey et al. 2008). This set of simulations allowed us to narrow down the values of *food* to a range that roughly corresponded to the patterns and values of larval survivorship (Bakker 1961; Mueller and Joshi 2000; Rodriguez 1989) and body size (Bakker 1961; Chiang and Hodson 1950; Rodriguez 1989) in *Drosophila* laboratory populations. The correspondence between simulation results and prior experimental data were judged by visual pattern-matching as per standard methodological recommendations (Railsback and Grimm 2011).

#### S4.2 Step 2. Calibrating the adult module

We next fine-tuned the parameters of the adult module, such that the fecundity of the female flies matched those of our flies (Dey et al. 2008). For the un-yeasted condition, we fixed *adnut* = 1, so that addens_ind_fec_i_ in the LHS of eq 4 represented the adult density-independent fecundity of females. We arbitrarily fixed *x_6_* = 2 and simulated eq 4 across a range of values for *size_adult*. This range for *size_adult* was obtained from the larval module (after the calibration described above) for different values of `numegg`. Our aim was to obtain a value for *x_5_* and range for *sen_adsize* that that led to a matching of the empirically observed qualitative patterns of addens_ind_fec with increasing *size_adult* (Chiang and Hodson 1950; Robertson 1957; Rodriguez 1989) (also see Text S3.2 above). We then calibrated the *sen_adden* parameter in eq 5 by visually matching the simulation results to the existing empirical patterns of adult density-dependent female fecundity decline (Fig 2 in Mueller and Huynh 1994, Fig 6.5 in Mueller and Joshi 2000).

At this stage, we had ranges of parameter values based on prior studies that roughly matched various life-history patterns from different prior single-generation experiments. We now used these ranges to calibrate the model for the various population dynamics indices for our experimental dataset.

#### S4.3 Calibrating for our dataset

Using the ranges of the various parameter values obtained above, we simulated 49-generation long time series experiments to fine tune the parameter values for *food*, *adnut*, *sen_adden* and *sen_adsize*. For this, we crossed the ranges of these parameter values to obtain a total of 129,920 parameter combinations (i.e. 8 values of *food* × 58 *sen_adden* × 20 *sen_adsize* × 14 *adnut*) and for each combination, we ran 50 replicate simulations. From these 50 replicate time series for each parameter combination, we computed the means of the average population size, *FI*, coefficient of variation (CV), and first and second autocorrelation lags of the simulation output. We then obtained the corresponding indices for the eight replicates each of LL and HH regimes of the experimental time series. The match between the simulation and experimental data was judged by computing the absolute of the percentage deviation between the simulation mean and experimental mean for each index and summing them up over the five indices. This led to a set of parameter values for which the sum over all deviations were minimized separately for the LL and the HH regimes. We then simulated the HH and LL time series regimes for these parameter values and did minor heuristic adjustments to obtain better matches (judged visually as per standard recommendations; Railsback and Grimm 2011) with the various indices of the dynamics of the experimental populations.

The above calibration process relied on fixing some parameters (e.g. *x_1_, x_2_* etc.) arbitrarily as constants, some based on prior experimental results (e.g. *m_c_*), and tuning the values of the other parameters to best match certain properties of the experimental data. Thus, the values of the best-fit parameters could be potentially very different if a different set of properties of the experimental data were chosen or the arbitrary constants were set to some other values. This is an inherent feature of pattern-oriented modelling. Since the main aim of this study was to vary these parameters and observe the corresponding effects on the dynamics, our calibration goal was not to arrive at impeccable parameter estimates but to derive a set of values that enabled our model to reasonably describe multiple facets of the empirical dynamics.

It should be noted here that once the values of *food* were obtained for the LL and HH regimes, we used the same values for the LH and HL regime. In other words, the LL and HH regime were equivalent to “training” datasets while the LH and HL regimes were equivalent to “prediction” datasets. This allowed us to avoid the issue of circularity in terms of parameterization and judging model performances. The final values of all the parameters used are presented in Table 1.

### Text S5. Comparisons with a previous model

Our model is similar to a previous model of the population dynamics of *Drosophila melanogaster* (Mueller 1988), with three major differences. First, the previous model was fully deterministic, while ours is stochastic and individual-based (for larvae and adults). This feature allowed us to study the various properties of the population size distributions and compare them with the experimental data, which would not have been possible with a deterministic model (except perhaps for chaotic dynamics). This also allowed us to account for certain biological features that can have a major impact on population dynamics. For example, allotting the sex of every individual using a Bernoulli distribution allowed us to account for demographic stochasticity in the number of females, even though the expected sex ratio was 1:1. The previous model, being deterministic, assumed that a fixed fraction of the individuals in the population were females.

Second, we considered female fecundity to be a logarithmic function of female body size, whereas in the previous study, it was modelled as a power law. This implies that in the previous study, when body size was large, small differences in size translated into large differences in fecundity, which was not the case with our model.

The third difference relates to the way effects of adult density on adult fecundity was modelled. In the previous model, low nutrition for adults refers to a condition where a dilute solution of yeast is provided (Mueller and Huynh 1994), as a result of which, yeast is not limiting at low densities and the fecundity of the flies is as high as the case when yeast is provided (see Fig 2 of Mueller and Huynh 1994). For this reason, in the previous model, the effects of adult nutrition are conceptualized solely as a density-dependent reduction of fecundity, assuming that when densities are low, the fecundities are the same. However, in our study, low food for adults means no yeast at all and consequently, the density-independent fecundity of our non-yeasted flies is much lower. That is why, we model the effects of adding yeast using the parameter *adnut* (which determines how much the fecundity is boosted due to yeast supplement) and assume that the density-dependent reduction in fecundity is similar in magnitude for both yeasted and non-yeasted flies. The latter condition is not a property of our model but gets imposed because we chose to use the same value of *sen_adden* for both LL and HH. Thus, this difference is simply a result of modelling strategies and the data used for model formulation and calibration.

Overall, as stated above, our model is more appropriately considered as a consequential extension of the earlier model (Mueller 1988), rather than a completely new model.

### Text S6. Role of extinction in the present study

Persistence, or the ability to resist extinction, is an important aspect of the dynamics of any population (Grimm and Wissel 1997). Prior empirical studies have suggested that extinction plays a crucial role in determining the dynamics of laboratory populations of *Drosophila melanogaster* (Dey and Joshi 2006; Dey and Joshi 2007). However, most of these studies have been conducted on the LH regime, under extremely low levels of larval food which leads to high extinction frequencies. For example, Dey and Joshi (Dey and Joshi 2006) provided ∼ 1mL of larval food and observed a per generation extinction probability of ∼0.37. However, the current empirical study used ∼2mL of food in the low food regimes, which led to a per generation extinction frequency of 0.06 (± 0.0067 SEM) in LH and 0.03 (± 0.01 SEM) in LL populations. The two other regimes did not suffer any extinctions at all. Therefore, there was no way for us to compare the regimes in terms of extinctions, which is why we chose to neglect extinction in this study, including in the process of model calibration. Interestingly, even though we neglected extinction during model calibration, the observed values of extinction frequency from the simulations for parameter values mentioned in Table 1, were fairly close to the experimental values (LH: 0.069 ± 0.013 SEM; LL: 0.003 ± 0.003 SEM; HL and HH: 0). We note here that these values refer to the extinction probabilities under parameter values that were close to our experimental conditions (i.e. Figure 1). When we studied the effects of changing the parameter values (Figures 2-6), it is entirely possible that the amount of extinctions observed were very different. Therefore, in principle, for every figure showing the effects of changing a parameter on constancy stability (here, *FI*), a corresponding graph on persistence stability (in the form of extinction probabilities or some related measure) can also be investigated. Although important in its own right, such an investigation is clearly out of scope of this paper which focuses primarily on dynamics in terms of constancy stability.

### Text 7. Model Recalibration for the dynamics of the FEJ-JB populations

To use the time-series data from an earlier experiment (Dey et al. 2008), we first re-calibrated our model by reducing the value of *m_c_* of FEJs from 1.1 to 1.0. This is because it has been suggested that due to selection for faster pre-adult development, the FEJs had a lower value of *m_c_* (Prasad et al. 2001). Moreover, to keep parity with the experimental data, we used 16 replicates each of FEJ and JB in both HL and LH nutritional regimes, and each replicate was simulated for 36 generations. Every other detail of the parameter values and the simulation were identical to those mentioned above. We then compared the population size distributions and *FI* values of the simulations against those observed from the empirical data

### Text S8. Simulating the dynamics of sex biased populations

We investigated the dynamics and stability of populations under three scenarios where there could be unequal numbers of males and females, namely systematic distortion of sex ratio, sex-specific culling and the sterile insect technique (SIT). In all these cases, relevant alterations were made in the adult module while the pre-adult module was kept unchanged.

a. Distortion of sex ratio: To simulate sex ratio distortions, we varied the probability of a given adult to be a female in the adult module, while keeping the rest of the model unchanged. We studied the effects of these various sex ratios at two different levels of density-independent basal fecundity, namely high [(*adnut* × *x_5_*) =1.49 × 85] and low [(*adnut* × *x_5_*) = 1.49 × 17)]. For each sex ratio × fecundity combination, we also studied the effects of low and high levels of food to the juvenile stage (*food* = 1.76 and 2.56 respectively). For each sex ratio × fecundity × food combination, we ran 100 replicate time series, each of which was 100 generations long. The mean (±SEM) of average population size and fluctuation index over these 100 replicates was plotted.
b. Sex-specific culling: This was similar to the above case except that in every generation, after the gender was assigned to the adults following strict 1:1 sex ratio, we explicitly removed a fixed percentage of males or females from the population. The culled individuals were chosen randomly. The effect of adult crowding on adult fecundity was computed after the culling.
c. Sterile insect technique (SIT): This is a popular pest control technique (Dyck et al. 2005), wherein large numbers of sterile males of the pest species are introduced into the population. These sterile males compete with the fertile ones for potential mating partners, thereby bringing down the number of successful matings which can lead to offspring. The number of females that mate with fertile males is directly proportional to the ratio of number of fertile: sterile males and inversely proportional to the competitive ability of the sterile males (Knipling 1955). For the sake of simplicity, we considered the proportion of females producing eggs for the next generation as a linear function of the fertile/sterile male ratio. Thus, proportion of egg laying females = min((`num_fertile`/(*β*×`num_sterile`)), 1), where `num_fertile` and `num_sterile` are the number of fertile and sterile males in the population, *β* is the competitive ability of the sterile males and min() is the minimum function. In this section, we studied the interaction of density-independent female fecundity with the amount of juvenile food available in determining the efficacy of SIT.

In our model, low values of *food* or high female fecundity can lead to extinctions due to larval overcrowding. This creates a problem in terms of judging the efficacy of introducing sterile males to cause extinction. Therefore, we first investigated the extinction probabilities over a large number of combinations of larval food (*food*, range = 0.5 - 3.0, step size = 0.2) and density-independent fecundity (i.e. *adnut* × *x_5_*, range = 1 - 150, step size = 5) in the absence of any introduced sterile males. For each *food* × fecundity combination, we computed the probability of extinction within 10 generations over 10 replicates and plotted the average extinction probability (Figure S10). We then limited our explorations on the efficacy of SIT to only those *food* × fecundity combinations where the probability of an extinction event within the first 10 generations was less than 0.1. For each *food* × fecundity combination, we sought to compute the minimum number of sterile males that would lead to an extinction within 10 generations. Therefore, we increased the number of sterile males in step size of 1 and simulated 10 generation long time-series for each *food* × fecundity × sterile male number combination till there was an extinction. For a given *food* × fecundity combination, this step was repeated 100 times and the average number of sterile males that were needed to cause extinction was recorded.

### Text S9. No evidence of transients in our experiments and simulations

Due to logistic constraints, most observed population dynamics time series tend to be short. However, it is well known that the transient behavior of population dynamics models can be very different from the equilibrium behaviors (Hastings 2004; Hastings and Higgins 1994). In this study, to keep parity with our experiments, we had limited the length of each simulated time series to 49. To investigate if the long-term behavior of these time series was any different from the short-term behavior, we simulated the dynamics in each regime for 1000 generations, and computed all the quantities represented in Figure 1 for the last 49 generations (Figure S6). Comparing generations 1-49 with generations 952-1000 revealed no major differences in either the population-size distributions or *FI*. This suggests that the transient dynamics in our model are almost indistinguishable from the longer-term dynamics.

Although their absence is re-assuring from a modelling perspective, transients are very much expected from a biological standpoint. This is because experimental evolution studies suggest that in *Drosophila melanogaster*, even 10-15 generations is often sufficient for noticeable divergence in life-history related traits that can affect the dynamics (for examples see Prasad and Joshi 2003). Therefore, all else being equal, one would expect at least some of the stability determining parameters to evolve during the course of the experiment, which in turn is expected to lead to transient dynamics in a long time-series. Yet, we did not incorporate any evolution in our model, which meant that the various life-history parameters detailed in Table 1, remained constant in a particular simulation run. This was because it has been previously shown that at least over 45 generations, there are no observable changes in stability determining demographic parameters in laboratory populations of *D. melanogaster* (Mueller et al. 2000). Thus, we felt that it was safe to ignore changes in life-history related parameters in the context of our empirical data and, therefore, did not incorporate their evolution in our model. However, we note that the structure of our model is such that it can be very easily extended to incorporate the evolution of stability-determining parameters and the effects of such evolutionary change on population dynamics.

---

**Figure S1.**
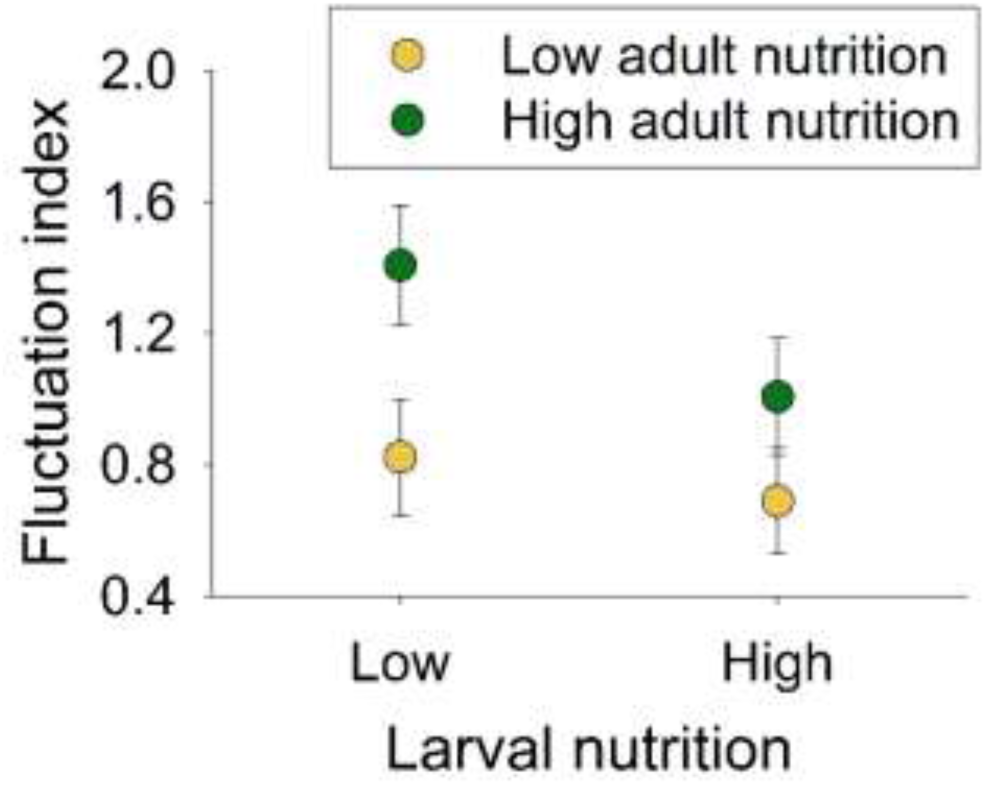
Effects of larval and adult nutrition on constancy stability. Interaction of larval and adult nutrition to determine constancy stability of population is statistically significant. High adult nutrition destabilizes population more when larval nutrition is low. Error bars = 95% CI.

**Figure S2.**
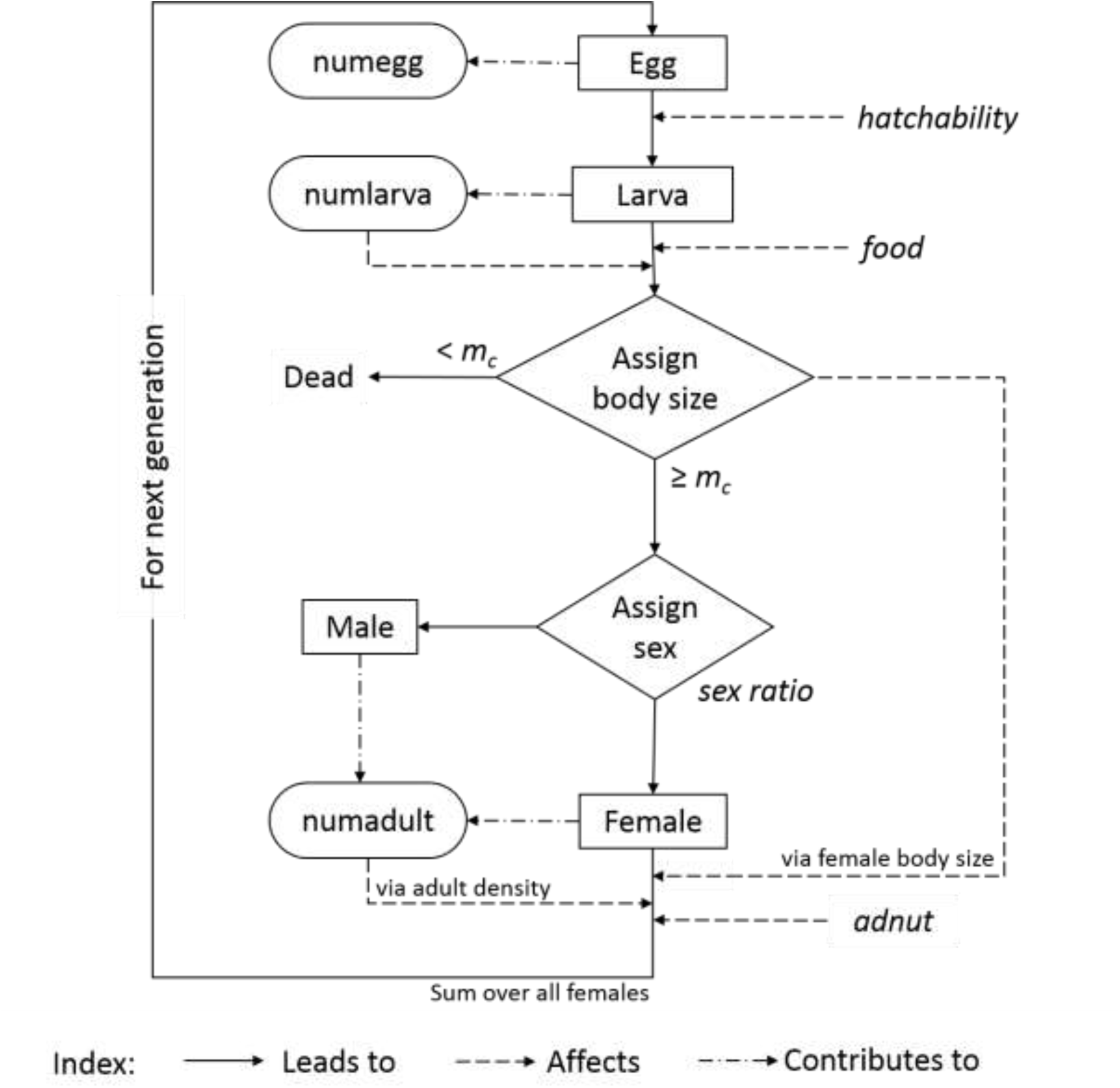
Schematic diagram of the model processes.

**Figure S3.**
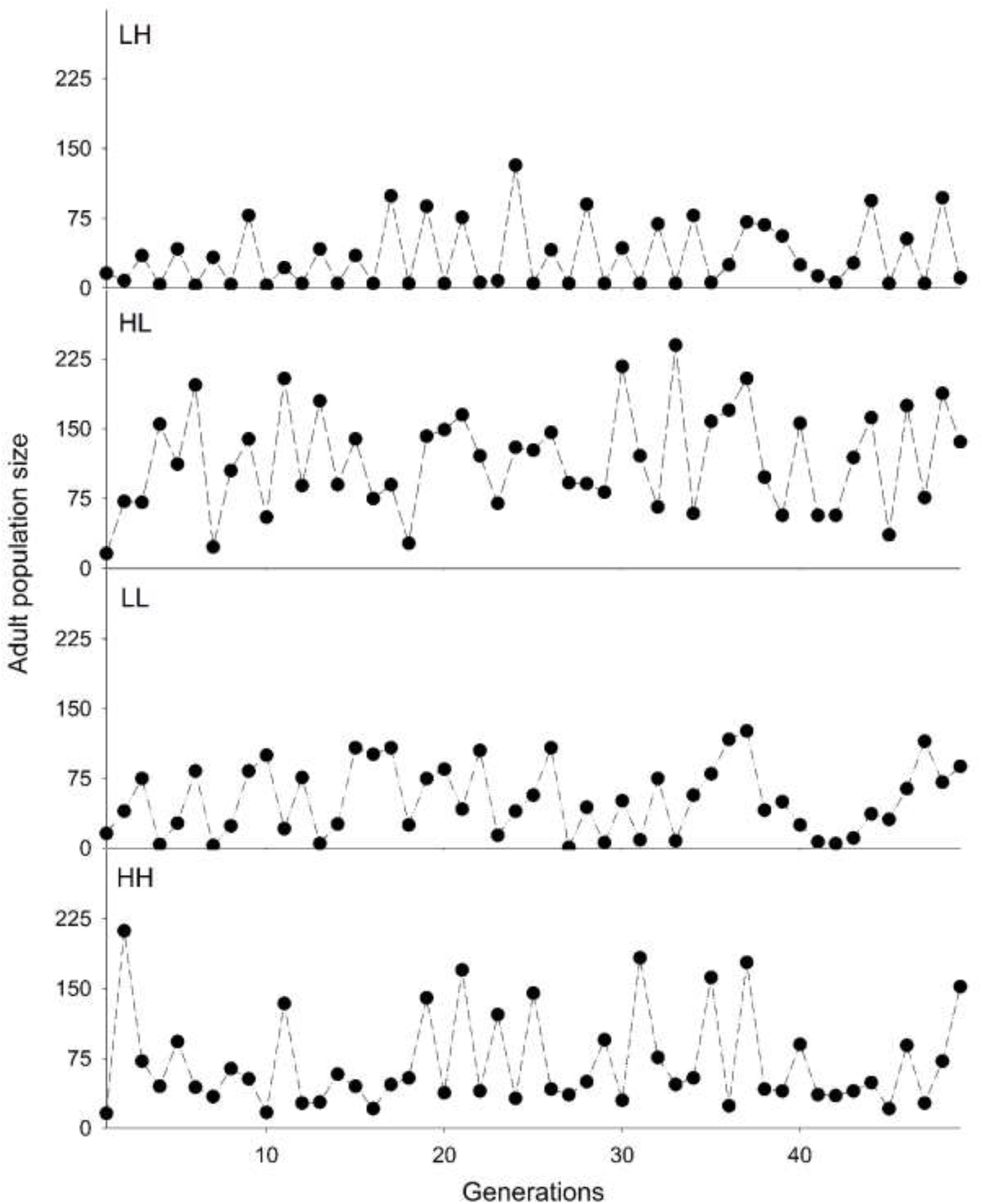
Representative empirical time-series of population size over 49-generations for LH, HL, LL and HH regimes respectively.

**Figure S4.**
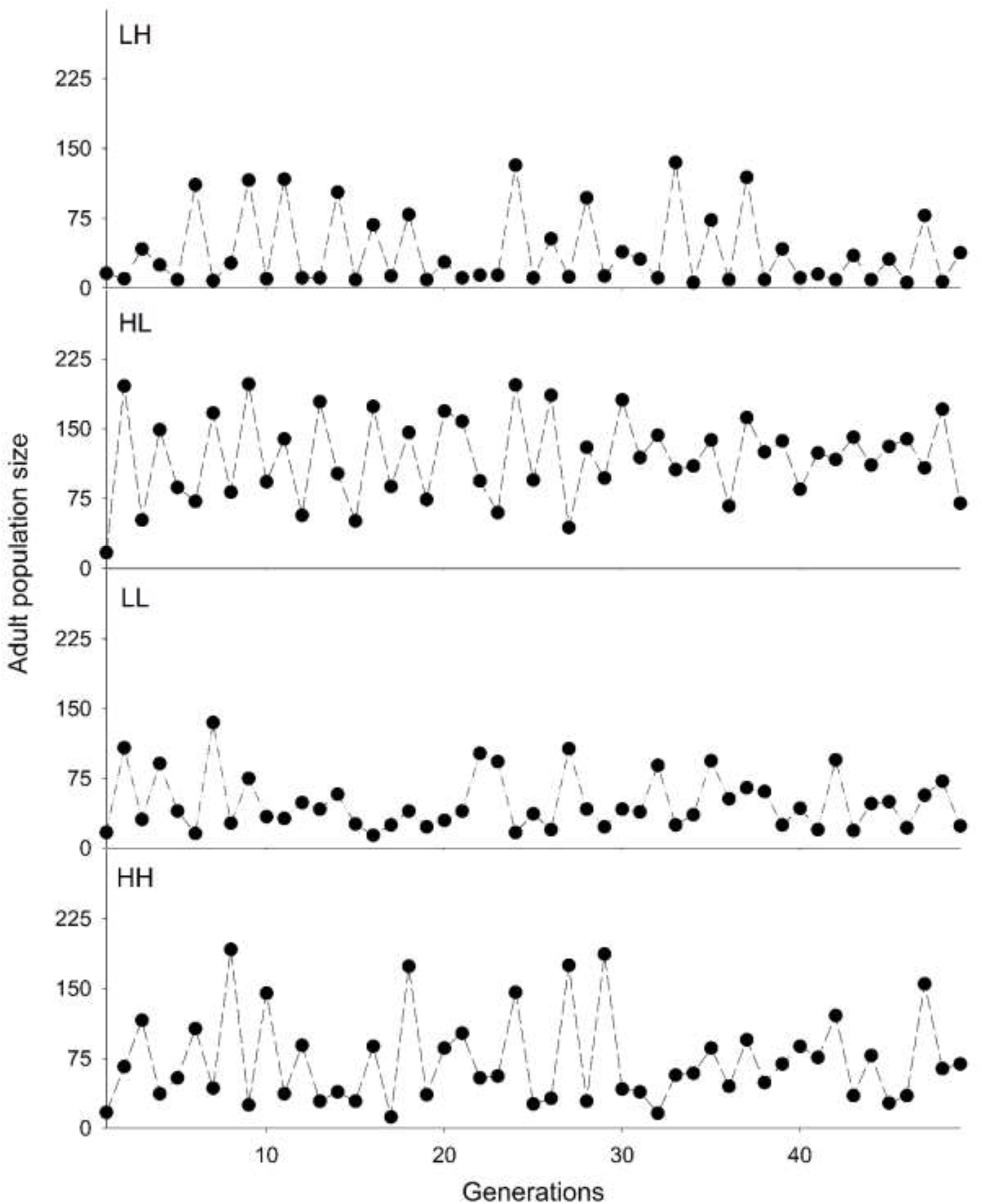
Representative simulation time-series of population size over 49-generations for LH, HL, LL and HH regimes respectively.

**Figure S5.**
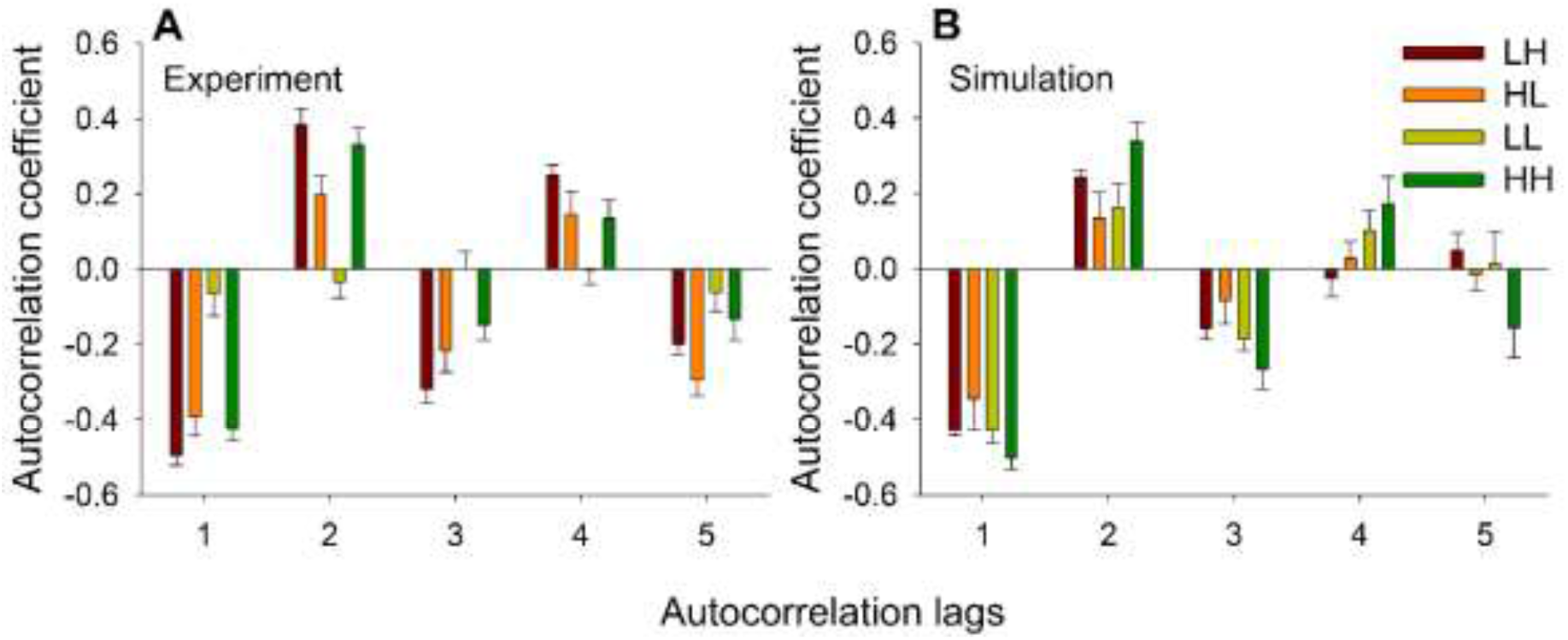
Autocorrelation coefficients of empirical and simulation time-series. For four regimes, we have computed first five autocorrelation lags, averaged over eight replicates for both (A) empirical and (B) simulation time-series. Error bars indicate SEM.

**Figure S6.**
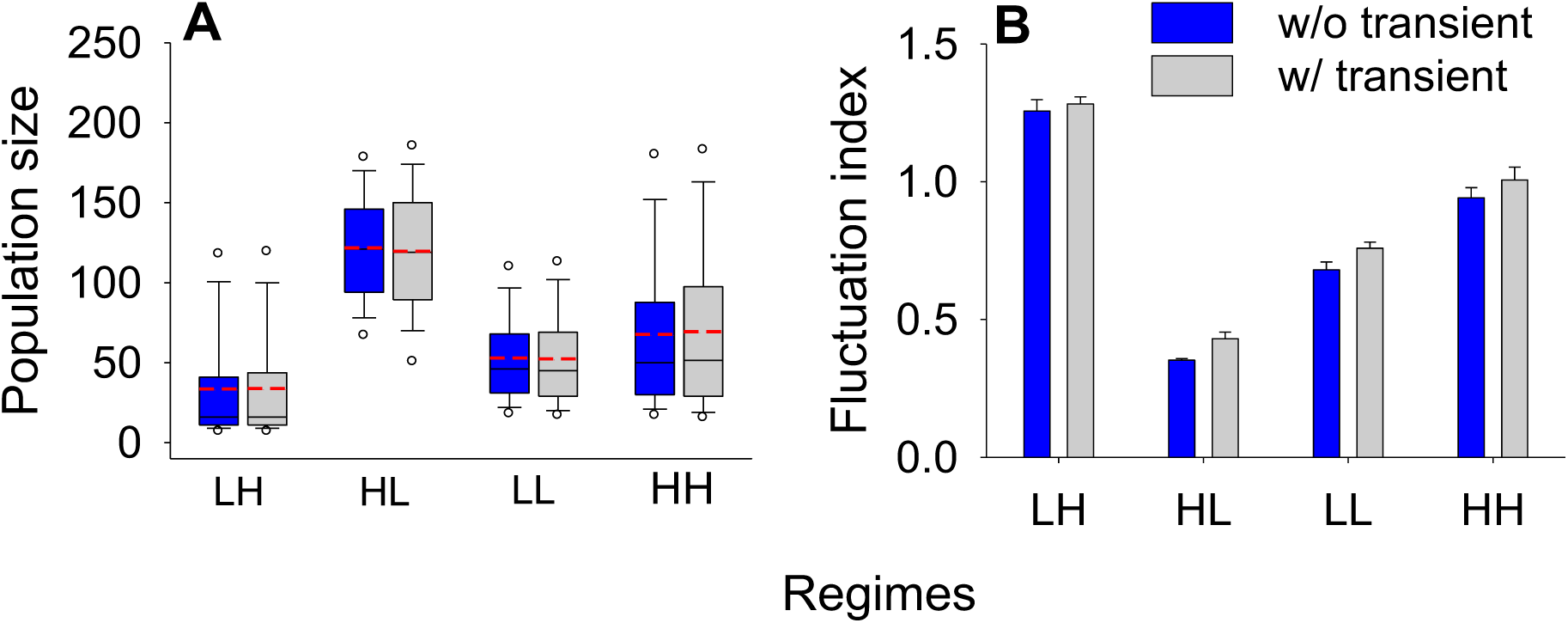
Population size distribution and constancy stability of short and long term dynamics. Blue boxes and bars represent the data for long term dynamics (generation 951-1000), where transients are excluded, whereas the grey shaded boxes and bars represents short term dynamics (generation 1-49). (A) Descriptive statistics of the population size distributions for long and short term dynamics in four regimes. Red dashed lines = means, thin black lines = medians, edges of the boxes=25^th^ and 75^th^ percentiles, whiskers=10^th^ and 90^th^ percentiles and the circles outside = 5^th^ and 95^th^ percentiles of the distributions. (B) Average (± SEM) fluctuation index of the population size distributions for long and short term dynamics in four regimes. In both panels, there are no systematic differences in population size distributions and constancy stability between the short- and long-term dynamics.

**Figure S7.**
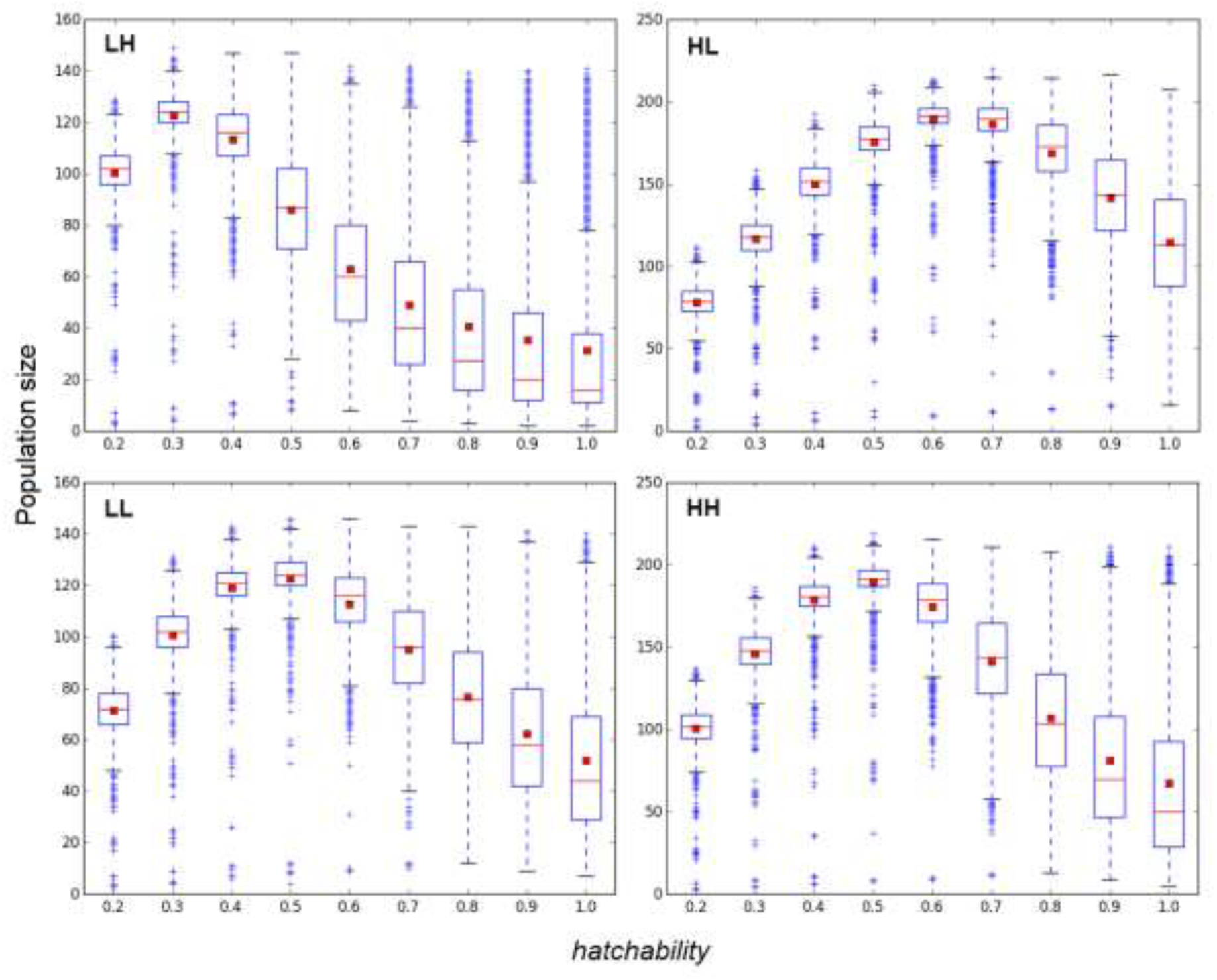
Effect of varying *hatchability* on population size distribution in four regimes. Red squares = means, thin red lines = medians, edges of the boxes=25th and 75th percentiles of the distributions.

**Figure S8.**
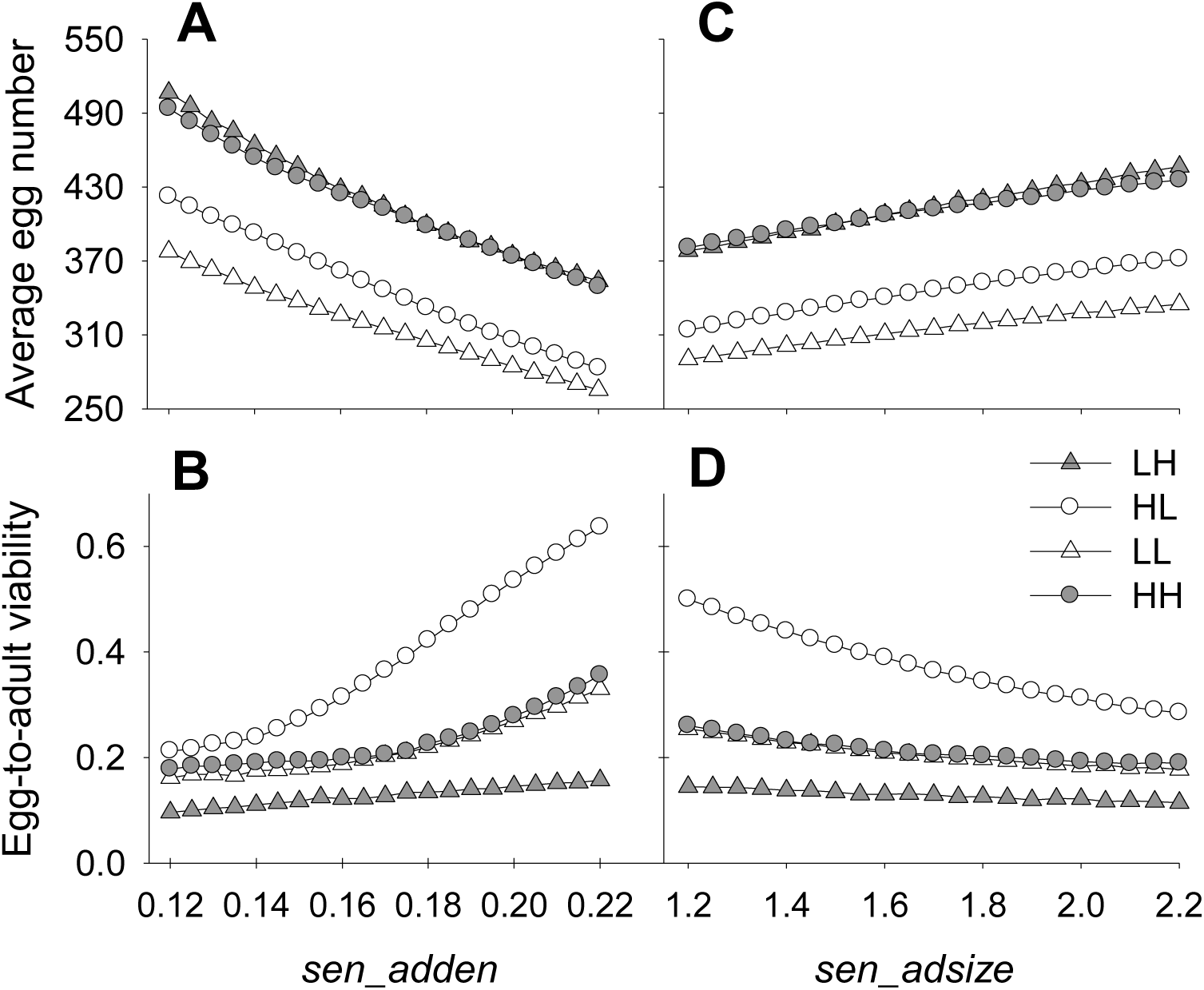
Effect of varying *sen_adden* and *sen_adsize* on the average egg number and egg-to-adult viability. Each point represents average (± SEM) fluctuation index of 100 replicates of 100-gen long simulated time series. Error bars are too small to be visible. Effects of sensitivity to adult density (*sen_adden*) on A. Average egg number and B. Egg-to-adult viability. Effects of sensitivity to adult size (*sen_adsize*) on C. Average egg number and D. Egg-to-adult viability. Note that in B and D LH is the least affected by increases in the parameter values. See text (section 3.4.2) for explanation.

**Figure S9.**
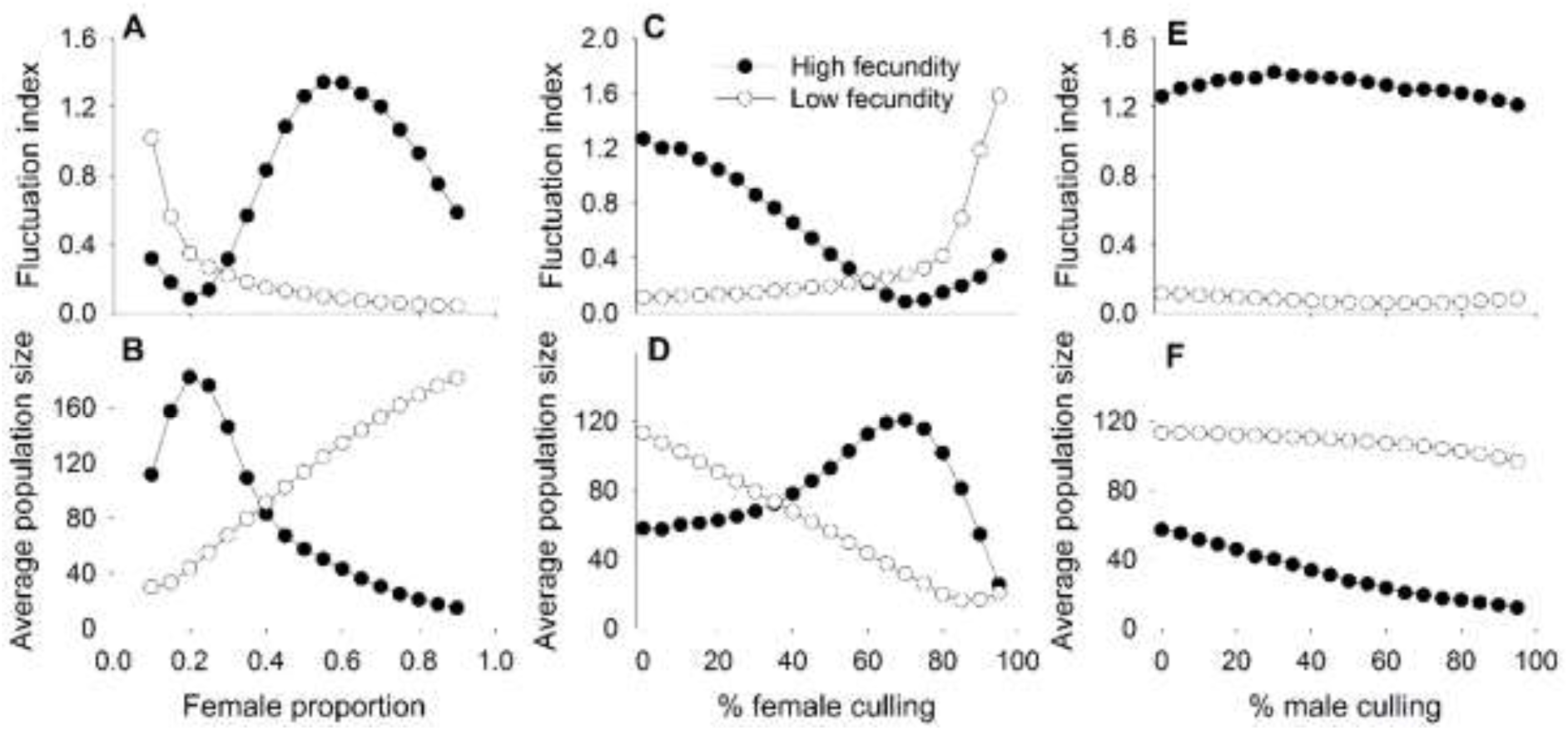
Effect of varying sex-ratio and sex-specific culling on population dynamics for high larval nutrient scenario. Fluctuation index and average population size were computed as the proportion of females (A and B), percentage of female culling (C and D) and male culling (E and F) increases in the population respectively. Each cases were investigated at two levels of density-independent fecundity. Here, each point represents average (± SEM) fluctuation index or population size of 100 replicates of 100-gen long simulated time series. Error bars are too small to be visible.

**Figure S10.**
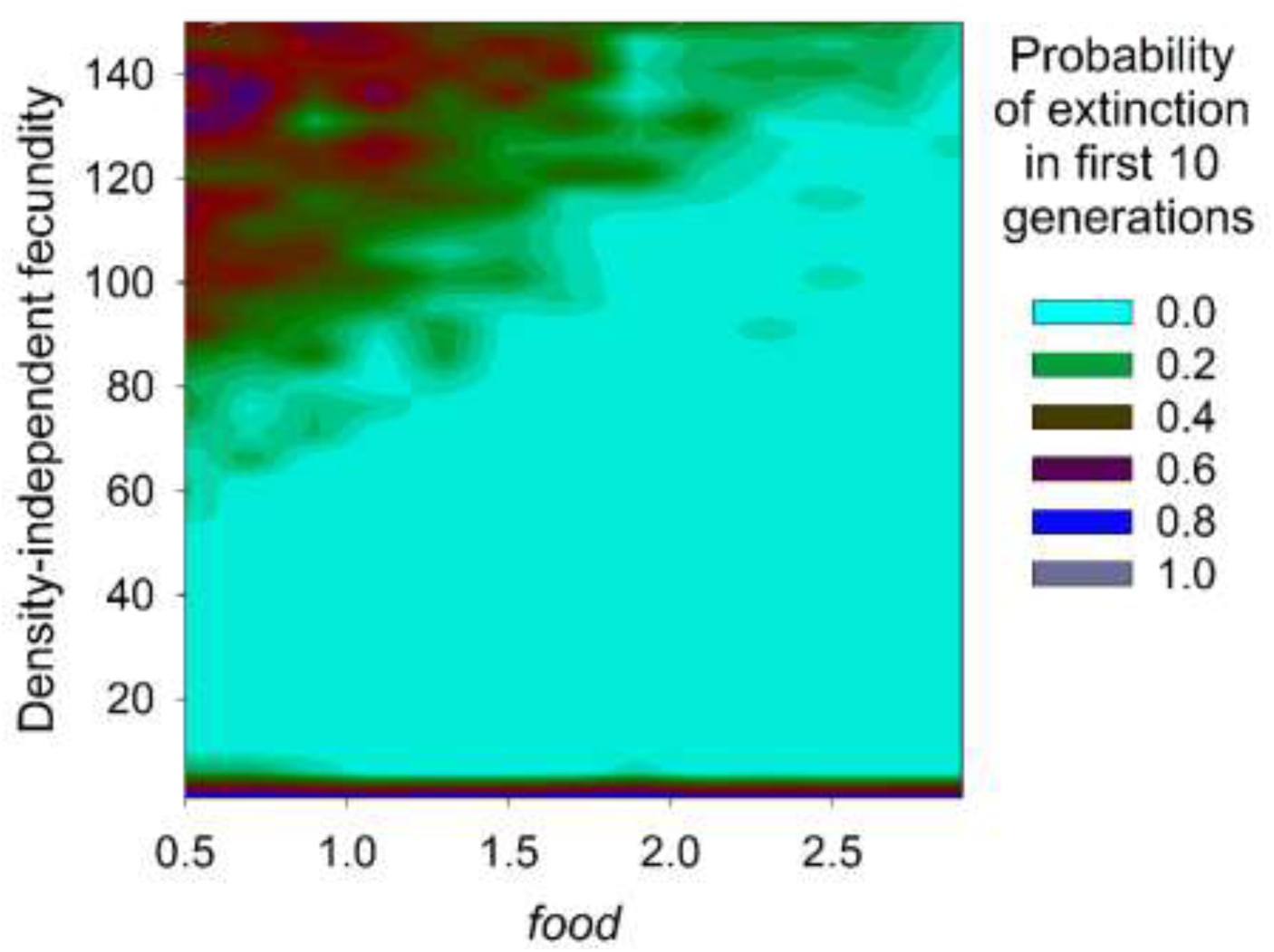
Probability of extinction within 10 generations in *food* × density-independent fecundity space. For each combination of food and fecundity values, we computed average probability of extinction within first 10 generations over 10 replicates and plotted according to the adjacent colour index.

